# Baltica: integrated splice junction usage analysis

**DOI:** 10.1101/2021.12.23.473966

**Authors:** Thiago Britto-Borges, Volker Boehm, Niels H. Gehring, Christoph Dieterich

**Affiliations:** Section of Bioinformatics and Systems Cardiology, Klaus Tschira Institute for Integrative Computational Cardiology, University Hospital Heidelberg, 69120, Heildeberg, Germany; Department of Internal Medicine III (Cardiology, Angiology, and Pneumology), University Hospital Heidelberg, 69120, State, Heidelberg, Germany; Institute for Genetics, University of Cologne, 50674, Cologne, Germany; Center of Molecular Medicine Cologne (CMMC)), University of Cologne, 50937, Cologne, Germany

**Keywords:** RNA-Seq, differential splicing, workflows, data integration, reproducibility

## Abstract

Alternative splicing is a tightly regulated co- and post-transcriptional process contributing to the transcriptome diversity observed in eukaryotes. Several methods for detecting differential junction usage (DJU) from RNA sequencing (RNA-seq) datasets exist. Yet, efforts to integrate the results from DJU methods are lacking. Here, we present Baltica, a framework that provides workflows for quality control, *de novo* transcriptome assembly with `StringTie2`, and currently 4 DJU methods: `rMATS, JunctionSeq, Majiq`, and `LeafCutter`. Baltica puts the results from different DJU methods into context by integrating the results at the junction level. We present Baltica using 2 datasets, one containing known artificial transcripts (SIRVs) and the second dataset of paired Illumina and Oxford Nanopore Technologies RNA-seq. The data integration allows the user to compare the performance of the tools and reveals that `JunctionSeq` outperforms the other methods, in terms of F1 score, for both datasets. Finally, we demonstrate for the first time that meta-classifiers trained on scores of multiple methods outperform classifiers trained on scores of a single method, emphasizing the application of our data integration approach for differential splicing identification. Baltica is available at https://github.com/dieterich-lab/Baltica under MIT license.

## Introduction

Alternative promoters, splice sites, and polyadenylation sites define the transcriptome complexity by producing different transcript isoforms. Alternative splicing (AS), defined as the differential removal of introns by alternative splice site usage instead of canonical splice sites, is widespread in eukaryotic genomes. AS regulation is central to physiological processes, such as tissue remodeling[1], and defective splicing has been linked to human disease[2]. However, most of the cataloged AS events are yet to be associated with their functional consequence[3]. Furthermore, there are increasing numbers of genomic single nucleotide variants associated with missplicing events[4], pointing to a latent link between multifactorial diseases and mis-splicing due to genetic variation. AS is regulated by context-dependent proteins named splicing regulatory factors, which define splice sites and lead transcript isoforms changes. Altogether, AS and its regulation are crucial to studying human health and disease. Computational methods for AS identification from RNA sequencing (RNA-seq) have helped to scale up these discoveries.

There are different approaches to identifying splicing events from RNA-seq. Methods that model intron usage are popular methods, as shown in Supplementary Figure S1. In addition, these methods have been applied to a broad range of studies, for example, the effects of genetic variation in splicing[5], identification of splicing factor-mediated AS events [6], associate AS to nonsense-mediated decay[7] and testing for splicing therapeutical intervention in animal models[8]. We here name these methods as different junction usage (DJU) methods. As suggested by Mehmood and collaborators[9], comparing results from multiple methods could improve AS event prioritization. Despite the popularity and critical application to human health, individual DJU methods have limitations.

DJU methods differ in software granularity. While some methods implement multiple functionality steps, from sequencing read filtering to results reporting, others focus solely on statistical modeling of RNA-seq split reads. These differences in implementation and poorly defined concepts are barriers to data integration from DJU method results. Specifically, DJU methods results are not comparable, as not all methods output standard file formats. Second, differences in AS event definition limits the comparison of event-specific metrics. The PSI (percent spliced in; Ψ) represents the proportion of splice site usage within an AS event per experimental group and indicates effect size[10]. In general, methods do not adopt a standardized definition for AS event or Ψ, thus complicating the comparison of effect sizes. Third, methods do not share common steps to facilitate result integration and benchmark. For example, it is not trivial to input the same matrix of splice junctions (SJ) read counts to all DJU methods. Collectively, these points are obstacles to data integration.

In this paper, we present Baltica, a framework that facilitates the execution and enables the integration of DJU methods results. Baltica comprises of a command-line interface, snakemake[11] workflows, containers[12], and scripts that provided reports on the integrated results. We propose a protocol to integrate results from DJU methods and further prioritize introns that undergo AS based on the decision of such methods. Optionally, Baltica integrates of results obtained with orthogonal experiments, such as AS evidence from Oxford Nanopore Technologies (ONT) RNA-seq. To our knowledge, there are no others solutions for integrating DJU results. We apply Baltica to 2 datasets. The first uses spike-ins with known experimental group concentration and transcriptome structure, the so-called Spike-in RNA Variant Control Mixes (SIRVs). The second ones uses paired Illumina and ONT RNA-seq datasets. In addition, Baltica integration allows us to compare the performance of different DJU methods and test the usability of a meta-classifier trained on the decision of the methods.

## Material and Methods

### Baltica method overview

Figure 1 shows an overview of the features included in the Baltica framework. Baltica comprises a commandline interface, workflow implementations, and scripts that handle DJU methods’ result parsing, integration, annotation, and reporting. The framework requires `snakemake`[11], and `singularity`[12]. Singularity containers and Bioconda[13] handle the dependencies for workflows and scripts. The containers allow the execution of software dependencies in isolation and provide reproducible workflows that don’t require direct user instalation.

**Fig. 1.**
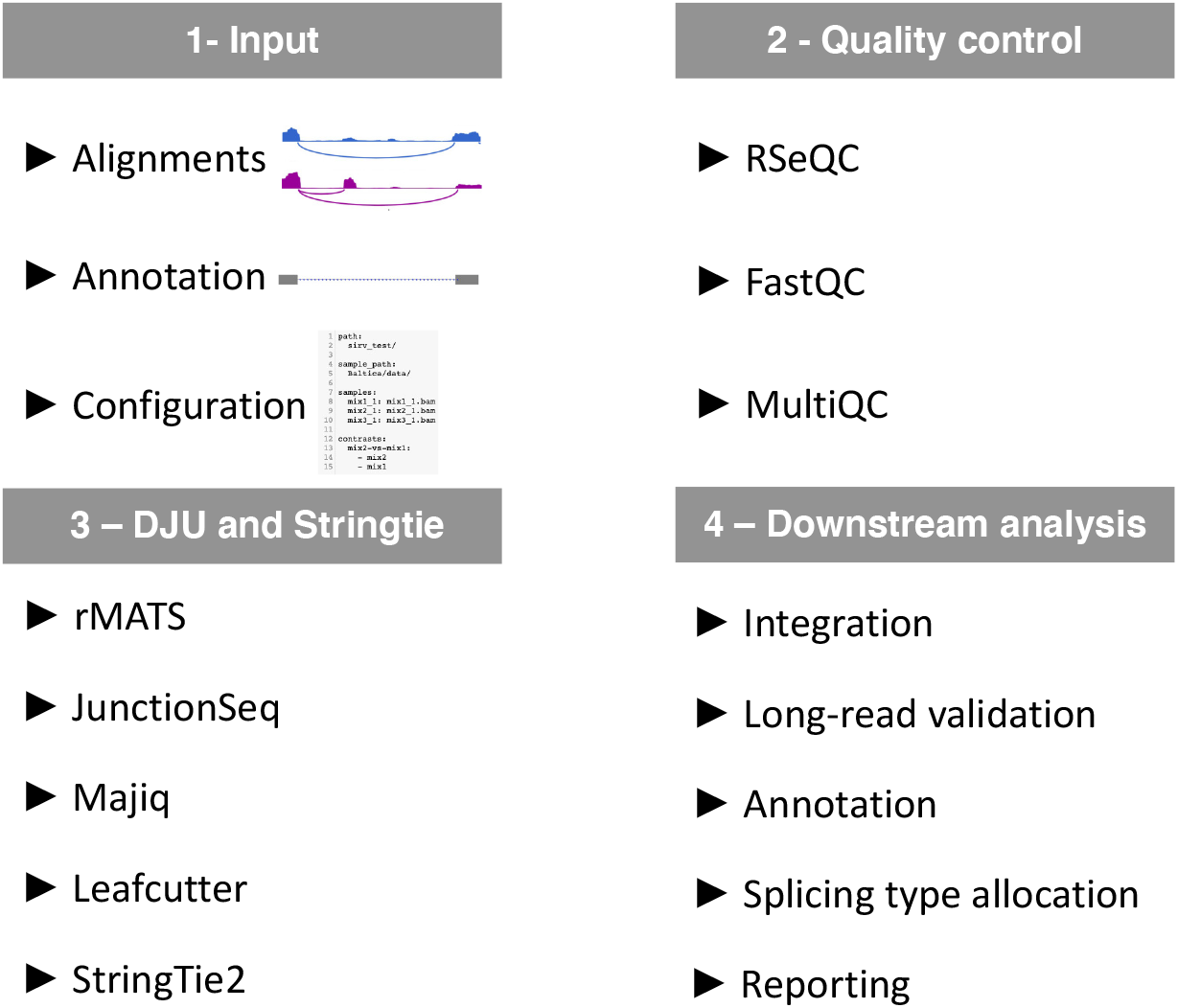
Baltica framework overview. Baltica is a framework to execute and integrate differential junction usage (DJU) analysis. **1 – Input:** Baltica takes as input RNA-seq alignments, reference annotation, and a configuration file. **2 – Quality control:** First, Baltica performs quality control of alignments with RSeQC and FastQC, which is reported by MultiQC. **3 – DJU and StringTie**: Next, Baltica computes DJU with rMATS, JunctionSeq, Majiq, and LeafCutter, and uses StringTie to detected transcripts and exons, which deviate from the reference annotation. **4 – Downstream analysis:** Finally, we integrate the results from the DJU method. Optionally, Baltica can include an extra piece of evidence for DJU (hereafter the orthogonal dataset), such as DJU obtained from Oxford Nanopore Technologies (ONT) RNA-seq. The set of introns is re-annotated using information from *de novo* transcriptome annotation, and splice types between SJ and exons are assigned. Finally, Baltica compiles a report with the most relevant information.

Baltica works as a standard Python package, and its command-line interface facilitates the execution of the workflows by, for example, automatically handling `singularity` arguments. The configuration file centralizes the required information for workflows. Specifically, it contains file paths, file to group assignments, method parameters, and pairwise comparisons between experimental groups to be tested. The required inputs are RNA-seq alignment files in BAM format, a reference transcriptome annotation (GTF/GFF format), and its sequence (FASTA format). Users can also input results from other evidence sources prepared in BED or GFF formats if available.

Baltica implements workflows for quality control methods, DJU methods, *de novo* transcriptome assembly, and downstream analysis. The included methods for quality control are `RSeQC`[14] and `FastQC`[15]. We use `MultiQC`[16] to summarize the output from both tools. In addition to the quality control of reads and alignments, this step helps to identify systematic differences among the analyzed samples or conditions. `RSeQC` implements an SJ saturation diagnostic, which quantifies the abundance of known and novel SJ. The tool also provides the proportion of reads per feature in the input annotation, which may indicate splicing changes due to, for example, an increase of reads mapping to introns.

Currently, the frameworks supports 4 DJU methods: `rMATS`[17], `JunctionSeq`[18], `Majiq`[19], and `LeafCutter`[20]. We detail the method inclusion criteria and workflows at Section 2.2. Finally, the analysis workflow proceeds with DJU integration, annotation, and reporting. Scripts for the analysis workflow were developed with R[21] and based on Bioconductor’s infrastructure to handle genomic coordinates[22, 23] as well as tools from the Tidyverse[24].

### Differential junction usage algorithms

Due to the high number of DJU methods available in the literature, we have established a set of rules for method inclusion into Baltica. We may include a method if it fits the following criteria:

- supports as input RNA-seq read alignment in the BAM format and transcriptome annotation in the GTF/GFF format
- provides test statistics, such as *p*-value, at the event or SJ level for pairwise comparisons
- outputs effect size estimates, such as the Ψ
- detects SJ independent of the reference annotation

We present an initial set of 4 DJU methods. These methods fulfill the criteria and are among the most popular methods for differential splicing identification, as shown in Supplementary Figure S1. But are we are aware that other DJU software packages exist, such as SUPPA2[25] and PSI-Sigma[26]. Therefore, we hope to include more of these packages into Baltica, especially with the help of the user community.

### rMATS-turbo

`rMATS-turbo` (v4.1.1), or simply, `rMATS`, estimates the splicingtype specific isoform proportion from RNA-seq reads. First, `rMATS` uses the reference annotation to determine the splicing events grouped by splicing types: skipped exon, mutually exclusive exons, alternative 3’ splice site, alternative 5’ splice site, retained intron. More recently, `rMATS`’ developers released experimental support for unannotated introns with the ‘−novelSS’ argument. `rMATS` uses the effective length-scaled junction read counts and, optionally, exon read counts to estimate Ψ. Then, it applies the likelihood-ratio to test whether ΔΨ (ΔΨ = Ψ_*i*1_ – Ψ_*i*2_, for the intron *i*, and groups 1 and 2) surpasses the 0.05 threshold.

### JunctionSeq

`JunctionSeq` (v1.16) takes as input a read count matrix obtained with `QoRTs`[27] (v1.1.8), for annotated SJ, novel SJ, and exons, so in fact, `JunctionSeq` falls into the differential exon usage and DJU classes. Based on `DEXSeq`, `JunctionSeq` uses disjoint genomic bins as features, and applies a generalized linear model[28] to model the feature expression. Beyond modeling the modeling aspect, `JunctionSeq` also invests in the visualization of the exon and intron usage and builds tracks for genome browsers. `JunctionSeq` does not identify splicing events, so the results are associated with intron coordinates.

### Majiq

`Majiq` (v2.2-e25c4ac) generates splice graphs for genes present on the RNA-seq dataset and the reference annotation. Next, it detects splicing events, quantifies the SJ usage from normalized SJ read counts, and computes the PSI value for the sample groups. `Majiq` uses a Bayesian framework to assess which ΔΨ changes threshold among groups are significant by a user-defined probability. The local splicing variations implementation includes more than 2 SJ per event. So it supports complex AS event types, which is more realistic than modeling splicing events by SJ pairs.

### LeafCutter

`LeafCutter` (v0.2.7) uses regtools[29] to extract and select reads SJ from RNA-seq alignments. Next, it uses an iterative clustering procedure to eliminate SJ with low usage. Finally, the `LeafCutter` fits a Dirichlet-multinomial generalized linear model on SJ usage proportion within intron-clusters.

A more detailed description of the workflow implementation is available at Baltica manual online[30].

## Baltica integration and reporting

To parse, integrate and annotate the results from the DJU methods, we use the Bioconductor infrastructure. While parsing the results files from the methods, Baltica pivots the results tables, so each row in the data table corresponds to a single SJ. Because `rMATS` outputs one result file for each AS event type, Baltica selects the SJ representing feature inclusion and exclusion events from each file. Because `LeafCutter` and `rMATS` assign the test statistics to the event instead of the SJ, we assign the same test statistics to multiple SJs contained in the AS event.

One challenge to integrating results from DJU methods is correcting for different coordinate systems. For example, methods can use 0-indexed (BED format) or 1-indexed (GTF format) files and use exon or intron splice site coordinates to represent the SJ genomic position. We make no assumptions regarding the method choice for the coordinate system, and this flexibility allows us to support many methods. To overcome the issue without fixing the coordinates adjusted for each method, we first compute the genomic overlap between introns in the reference annotation (subject) and a set of SJ output from a method (query). Then, we compute coordinate offsets (in nucleotides) between subject and query, determine the most frequent difference in start and end coordinates between the 2 sets, and finally, apply corrections to the coordinates in a strand-specific manner for each method. This procedure allows Baltica to report groups of SJ that represent splicing events in different genomic coordinate systems.

Next, Baltica uses a *de novo* and guided transcriptome annotation as a reference for annotation and assigning alternative splicing types. The de *novo* workflow comprises merging the alignment files for experimental groups; next, we use `StringTie` (v2.1.5)[31] to obtain group-specific transcriptome annotations, which are then subsequently combined with `gffcompare`[32] in the guided mode. We use this novel annotation for downstream analysis, including naming genes and transcripts and assigning AS types when possible. Novel SJ, not included in the reference transcript annotation, are also annotated. Currently, Baltica determines the following types: exon skipping (ES), alternative 3’ splicesite (A3SS), alternative 5’ splice-site (A5SS). The AS type assignment procedure occurs by comparing SJ to overlapping exons features, detected in the *de novo* annotation. We can determine the AS type using distance rules between the start and end coordinates of the SJ and its overlapping exons. Associating the SJ to transcripts enables the study of the splicing event in the context of the transcript sequence and structure. Finally, the framework produces a report that summarizes integration results. It provides an overview of the integration results and an HTML table with one SJ per row, the methods score, SJ annotation, and link to the UCSC Genome Browser[33].

## Benchmark

Methods to detect DJU from RNA-seq are valuable tools for prioritizing mechanisms driving splicing changes. From a classification perspective, differential splicing methods aim to classify introns that are truly differently used from the other introns. We approach the differential splicing identification as a binary classification problem. Thus, the positive instance, the differently spliced intron, is more relevant than the negative instance. For the SIRV dataset, introns in the SIRV transcriptome that have fold-change = were considered,positive,while others introns that were not changing (foldchange of 1) were negative instances. For the paired Illumina-ONT RNA-seq dataset, introns with *p*-value <0.05 were considered positive instances. This value was obtained with the `edgeR::diffSpliceDGE` function. The set of introns from the SIRV or ONT RNA-seq dataset were used as a reference, so the results among methods are comparable. A true-positive instance (TP) was defined as truly changing and correctly classified, while false-positive (FP) was a negative instance classified as positive. Accordingly, true negatives (TN) and false-negatives (FN) were true negative instances that were correctly or incorrectly called, respectively. The following metrics are defined:

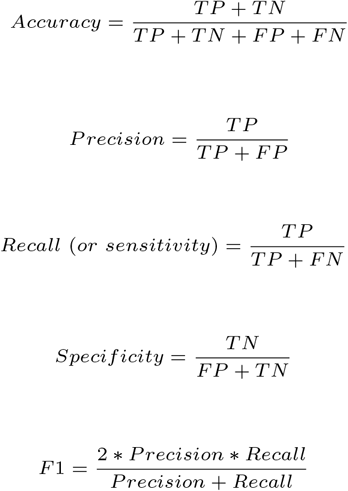

True positive rate (TPR) and false positive rate (FPR) are synonymous to recall and 1— specificity, respectively. Receiver Operating Characteristic (ROC) curve, Precision-Recall (PR) curve, and area under curve (AUC) were computed with ROCR[34]. Confusion matrix and associated statistics report were computed with caret[35], and for that, method scores were made binary using the 0.95 threshold. Heatmap and UpSet plots were created with ComplexHeatmap package[36].

### Meta-classifier to identify differential splicing

We propose a machine learning approach for a meta-classifier that combines the score of the DJU methods workflows implemented in Baltica. To do so, we train the models with the matrix of DJU scores from the second dataset, with matched third-generation sequencing. The dataset was split into training and testing data (80% vs 20%). DJU scores from the Illumina and ONT RNA-seq were used as input and target values, respectively. Feature selection proceeded with the mlxtend package[37] using either a method score column or a combination of the 4 score columns. Next, a grid search was performed with parameters for the Gradient Boosting Classifier (GBC) and Logistic Regression Classifier (LRC) algorithms implemented in scikit-learn (v0.24.2)[38], for features listed in Table 2, and the combination of columns. The grid search aimed to maximize the area under the ROC curve. This experiment allows us to compare the classification performance from the meta-classifier and classifiers trained from a single method score.

### RNA libraries preparation, sequencing, and alignment for the SIRV dataset

Figure 2 schematizes the application of Baltica to the SIRV dataset. Figure 2b compares DJU calls by the 4 methods and the ground-truth. Cell lines, RNA extraction, and RNA-seq were described in Gerbracht and collaborators[39]. In short, we obtained 15 libraries from Flp-In T-REx 293 cells, extracted the RNA fraction with TrueSeq Stranded Total RNA kit (Illumina), followed by ribosomal RNA depletion, with RiboGold Plus kit and Spike-In RNA Variants (Lexogen SIRV, Set-1, Iso Mix E0, E1 and E2, cat 025.031) input. Libraries were sequenced with an Illumina HiSeq4000 sequencer using PE 100bp protocol, which yielded around 50 million reads per sample. Data were deposited in ArrayExpress (E-MTAB-8461). Sequenced reads’ adapters and low-quality bases were trimmed, and reads mapping to human precursor ribosomal RNA were discarded. The remaining reads aligned with the human genome (version 38, EnsEMBL 90) extended with the SIRV annotation. In the DJU method benchmarking context, we are not interested in the actual biological condition but the SIRV transcriptome changes. Our experimental design does not confound a SIRV mix with the biological conditions, as detailed in Supplementary Table S1, so any AS events identified within human chromosomes are false calls. The SIRV transcriptome comprises seven genes, 101 transcripts, 138 unique introns, of which 98 change among the 3 mixes, leading to 294 changing introns. In conclusion, the 3 SIRV mixtures in the context of the complex human transcriptome allow us to compare the performance of the DJU methods.

**Fig. 2.**
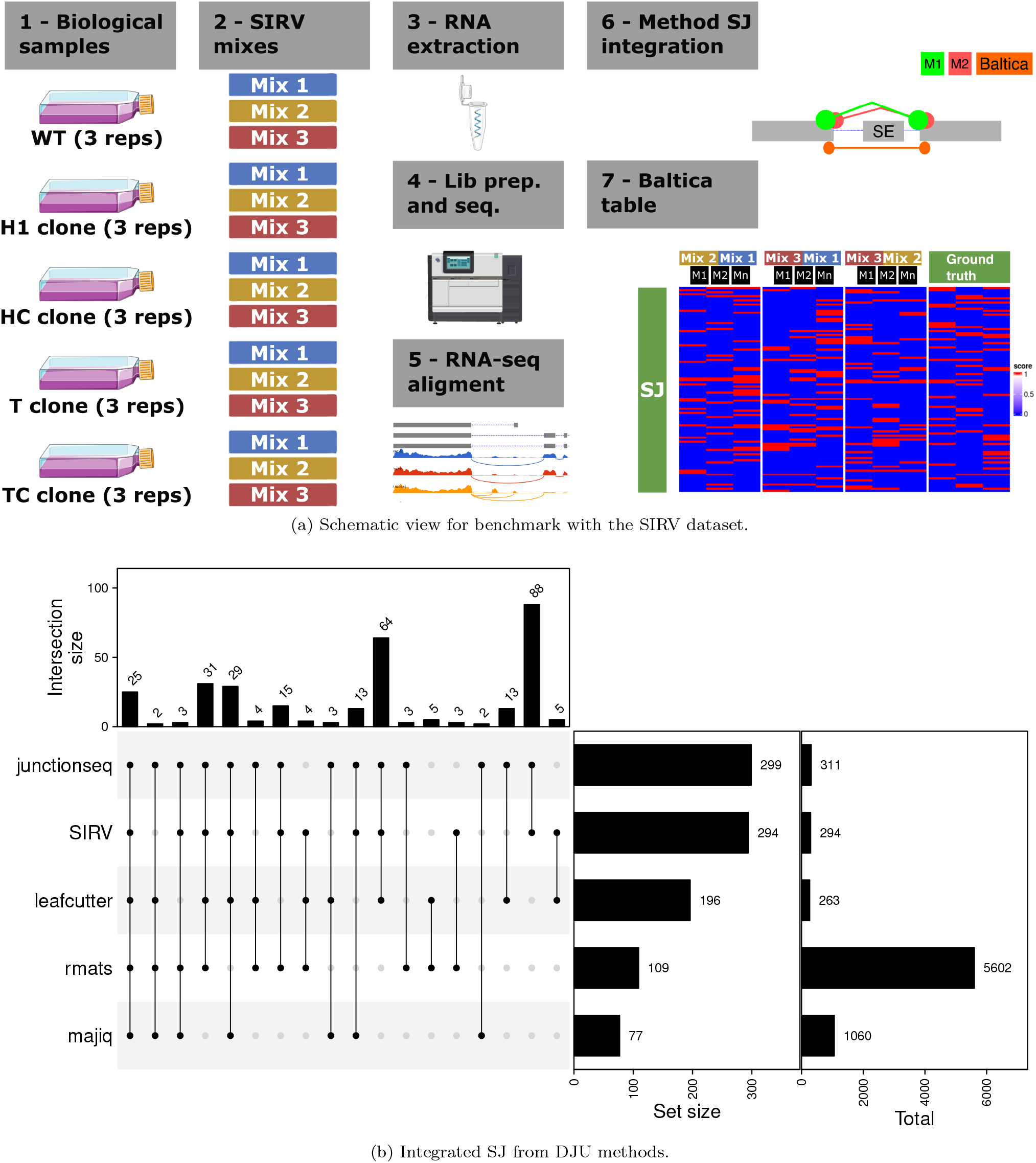
Integrated DJU results for the SIRV dataset. (a) The experimental design from Gerbracht et al.[39] has five biological groups in replicates, and Table S1 matches the biological samples groups to SIRV mixes and samples identifiers. SIRV mixes were included in a design not confounded to the biological groups. As detailed in Section 2.6, after RNA extraction, library preparation, and sequencing, the sequencing reads were aligned to the human genome extended with the SIRV genome. We apply Baltica workflows, as described in Section 2.1. To integrate the results, we first split AS events into individual SJ that are contained in each event. Next, we correct the start and end coordinates from SJ of multiple methods. Once SJ were integrated, we observed that the statistically significant SJ for `JunctionSeq` (padjust < 0.05), `LeafCutter` (p.adjust < 0.05), `Majiq` (probability_non_changing < 0.05) and `rMATS` (FDR < 0.05) have limited overlap with SJ that are known to change in the SIRV transcriptome. The score is defined as 1 — *padjust*, where padjust is the metric for the statistical test from each metric. In the figure, *M*_1_, *M*_2_,…, *M_n_* represent the multiple DJU methods. The UpSet plot in (b) shows distinct sets of introns called significant by combinations of methods and the SIRV annotation (294 true positive SJ). The intersection and set sizes show hits to annotated SIRV introns, while the total column shows the size of hits to the combined human transcriptome and SIRV annotation. The SIRV transcriptome has 98 distinct introns that change in fold change among the mixes. We omit the complement set for the combinations.

### RNA libraries preparation, sequencing, and alignment for the matched ONT RNA-seq and Illumina RNA-seq datasets

The cell lines, RNA extraction, library preparation, and RNA-seq have been described in Boehm et al. (2021)[7]. In detail, wild type (WT) or SMG7 knockout (KO) Flp-In-T-REx-293 cells were seeded on 2x 10 cm plates in high-glucose, GlutaMAX DMEM (Gibco) supplemented with 9% fetal bovine serum (Gibco) and 1x Penicillin Streptomycin (Gibco) at a density of 2.5xl06 cells per plate and reverse transfected using 6.25 *μ*l Lipofectamine RNAiMAX and 150 pmol of the respective siRNA (Luciferase as control for WT, SMG6 for SMG7 KO cells) according to the manufacturer’s instructions. Cells were harvested after 72 h with 2 ml of peqGOLD TriFast (VWR Peqlab) per plate and total RNA was isolated following the manufacturer’s instructions. The following changes were made: Instead of 200 *μ*l chloroform, 150 *μ*l 1-Bromo-3-chloropropane (Molecular Research Center, Inc.) was used. RNA was resuspended in 40 *μ*l RNase-free water. 100 *μ*g of total RNA was subjected to 2 rounds of consecutive poly(A)-enrichment by using 200 *μ*l Dynabeads Oligo (dT)25 and following the manufacturer’s instructions. Poly(A)-enriched RNA was eluted with 22 *μ*l RNase-free water and subsequently used for library preparation and ONT direct RNA sequencing. A total of 4 replicates per condition were sequenced. Reads were aligned with minimap2 (v2.17)[40]. We assembled the Nanopore reads and estimated junction read counts based on this new assembly with Stringtie2 (v2.1.1) and Ballgown (v2.14.0)[41]. DJU calls were computed with with EdgeR (v3.24.0)[42] using the diffSpliceDGE function. We use 1– *p*-value as the DJU score.

## Results

### DJU method performance comparison

We present benchmark experiments comprising the 4 DJU methods and 2 datasets, the SIRV dataset, schematized in Figure 2, and the paired Illumina-ONT-seq dataset.

### Benchmarking the SIRV dataset

The SIRV dataset comprises 414 SJ from the 3 comparisons with the 3 SIRV mixes. The benchmark shows that `JunctionSeq` outperforms all of the other methods (Figure 3). `JunctionSeq` ranks top with an AUC of 0.87, followed by `LeafCutter` (AUC 0.60), `Majiq` (AUC 0.59), and `rMATS` (AUC 0.53). While `JunctionSeq` performs best, `rMATS` performs close to a random classifier. Supplementary Figure S3 shows the PR curve, which shows the trade-off between precision and recall independent of method score. Of note, `Majiq` presents a low score variability.

**Fig. 3.**
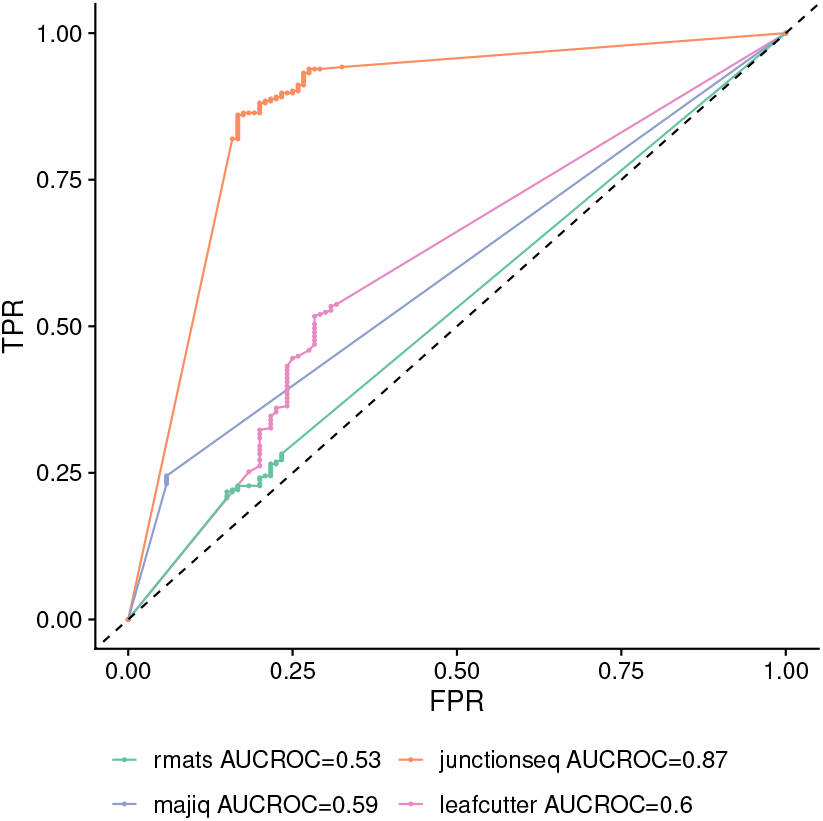
Baltica benchmark for DJU methods with the SIRV dataset. The introns matching the SIRV transcriptome validated on whether these introns change or not in a given comparison. The performance rank for both curves is consistent between the ROC and PR curves (Supplementary Figure 3): `JunctionSeq` is the top ranking method, followed by `LeafCutter`, `Majiq`, and `rMATS`. AUC: area under the curve; PR: precion-recall; ROC: Receiver Operating Characteristic; TPR: True Positive Rate; FPR: False Positive Rate.

All methods have a specificity ≥ 0.68, which represents a fair predictive performance for negative instances (Supplementary Figure S2 and confusion matrices in Supplementary Section S-X.6). Methods’ scores show only a limited agreement across each other (Supplementary Figure S4).

### Benchmarking ONT direct RNA-seq dataset as validation

To complement the size limitation of the SIRV dataset, we also benchmark an alternative dataset. Different to the SIRV dataset, this is not a *bona fide* ground-truth dataset. However, we observe similar pattern for the ROC curves of both benchmarks (Figure 3 and Figure 4). Interestingly, we note a reduction of the differences in AUCROC among methods in the second dataset (Figure 4). Most notably, `rMATS` AUCROC increased from 0.53 to 0.65. In terms of AUCPR metric, `JunctionSeq` ranks first with AUCPR of 0.72, followed by `Majiq` (0.61), `LeafCutter` (0.57) and `rMATS` (0.48) (Supplementary Figure S6). Out of 20,744 introns labeled as positive, 21% were called by all 4 methods, and 3 out 4 methods already call 43% (Figure S5).

**Fig. 4.**
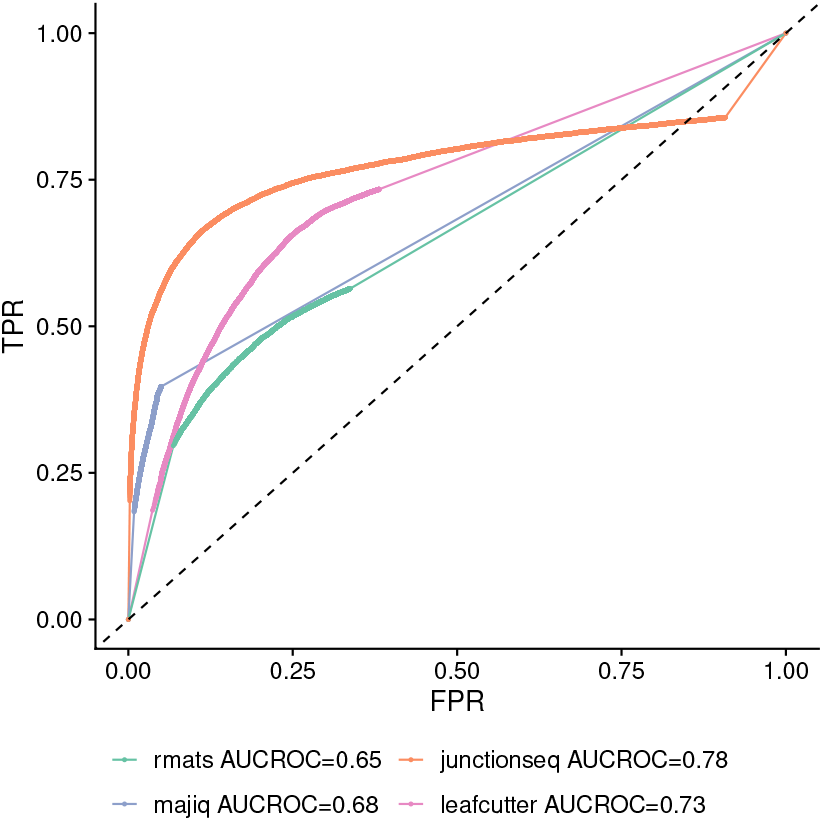
Baltica benchmark with a paired Illumina-ONT RNA-seq datasets. Performance for DJU methods executed with Illumina RNA-seq and validated with Nanopore RNA-seq show similarities with the benchmark with the SIRV dataset. The classification performance ranks are the same as the benchmark with the previous dataset, in Figure 3. `JunctionSeq` outperforms the other methods in both AUCROC and AUCPR (Supplementary Figure S6). However, in this case, the performance difference among methods is smaller than the benchmark with the SIRV dataset.

Table 1 compares the recall, specificity, and F1 metrics for the 2 benchmarks. There are multiple factors to explain the differences between the 2 benchmarks. Of note, the ratio of positive and negative instances change from 2/3 to 1/4 between the SIRV and paired Illumina-ONT RNA-seq datasets. This change can partially explain the overall gain in F1 for all methods, but `JunctionSeq`.

**Table 1.** Prediction metrics for the benchmark in the SIRV and ONT RNA-seq datasets. See the complete report at the Supplementary Section S-X.6 (SIRV) and S-X.11 (Illumina-ONT RNA-seq). Spec: Specificity.

|  | SIRV |  |  | Illumina-ONT RNA-seq |  |  |
| --- | --- | --- | --- | --- | --- | --- |
|  | Recall | Spec. | F1 | Recall | Spec. | F1 |
| rMATS | 0.26 | 0.77 | 0.39 | 0.51 | 0.76 | 0.61 |
| JunctionSeq | 0.91 | 0.74 | 0.82 | 0.65 | 0.90 | 0.75 |
| Majiq | 0.24 | 0.94 | 0.39 | 0.35 | 0.96 | 0.51 |
| LeafCutter | 0.54 | 0.68 | 0.60 | 0.70 | 0.70 | 0.70 |

**Table 2.** Parameter space for the grid search procedure with Gradient Boosting Classifier (GBC) and Logistic Regression Classifier (LRC). The procedure aims to maximize the ROC AUC(Area under the Receiver Operating Characteristic Curve) metricof scikit-learn’s implementation of the machine learning algorithms.In addition to these parameters, we compare classifiers trained in a single method versus a classifier trained on the 4 methods to test whether the differential splicing identification task benefits from a meta classifier.

| GBC Parameter | Parameter space |
| --- | --- |
| learning_rate | 1, 0.5, 0.25, 0.1, 0.05, 0.01 |
| max_depth | 1, 2, 4, 8 |
| subsample | 0.5, 0.8, 1.0 |
| min_samples_split | 0.2, 0.4, 0.8 |
| n_estimators | 1, 2, 4, 8, 16, 32, 64, 100, 200 |
| LRC Parameter | Parameter space |
| penalty | l1, l2 |
| C | $1.0 \times 10^{-3}$, $5.6 \times 10^{-2}$, 3.1, $1.7 \times 10^2$, $1.0 \times 10^4$ |

Methods reach a consensus in terms of classification of negative instances. In contrast, for positive instances, the consensus is not as clear (Supplementary Figure S7), and methods scores complement each other. The correlation among scores was low, with a maximum of 0.55 for the `LeafCutter` and `rMATS` pair (Supplementary Figure S8). The relatively low correlation and complementary nature of method scores for positive instances have motivated us to test the performance of a meta-classifier for differential splicing identification.

### Meta-classifier with combined DJU scores

Meta-classifiers may improve classification performance by combining the decision of multiple classifiers. To test the application of meta-classifiers to the differential splicing identification task, we have trained 2 models, the LRC and GBC, by fitting them with either a single feature (one method score) or 4 features, one for each method score.

Because models trained in a different set of features may require different sets of parameters, we apply a grid search over the parameters listed in Table 2. We observe the predictors with the combined scores outperform models fitted with scores from a single method score, independent of other parameters and the machine learning algorithm. Table 3 compares these results. To our surprise, the LRC algorithm performs competitively with the GBC algorithm. The GBC algorithm scores an AUCROC of 0.92 in the training set and 0.91 in the testing set, confirming the model’s ability to generalize to unseen data. This result demonstrates that combining scores from multiple DJU methods is favorable for differential splicing identification. It improves the predictive performance of the classifier targeting class determined by an orthogonal set, the ONT RNA-seq.

**Table 3.**
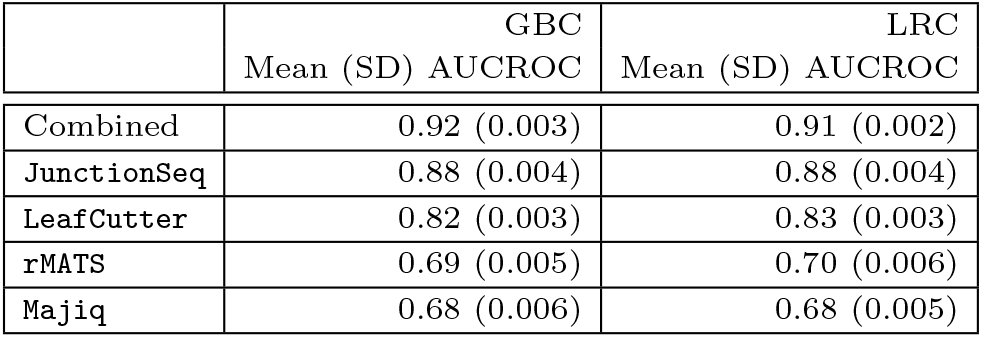
Comparison of the top-scoring meta-classifiers performance. Mean (standard deviation) AUCROC for the 10fold cross-validation. Each row represents the top-scoring model for models trained on the combined features of a single feature. SD: Standard deviation.

## Discussion

Baltica aims to enable the study of integrated results from DJU methods. To achieve that goal, the framework provides workflows[11] and containers[12] to execute the said methods. Next, it combines and re-annotates the results and reports them as an interactive table, as illustrated in Figure 5.

**Fig. 5.**
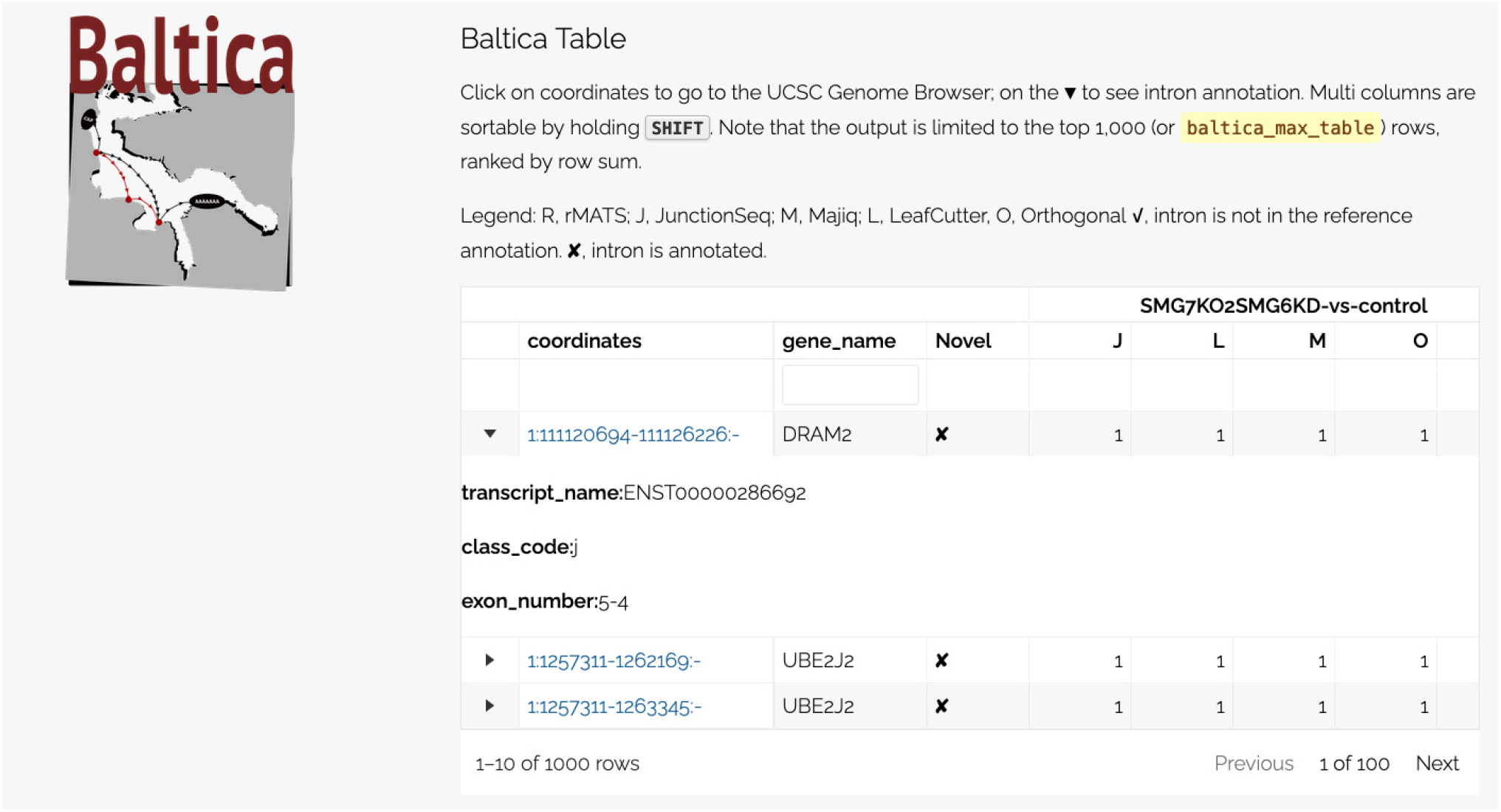
Interative Baltica table. The image shows a static view of the Baltica report, focusing on the table component. The table is pre-sorted by sums of the scores columns, and it can be sorted by the scores or filtered by gene name. In addition, annotation is provided for introns that match the de novo transcriptome annotation. Introns that are annotated are flagged in the **Novel** column. **J**, **L**, **M**, **R** and **O** stand for JunctionSeq, LeafCutter, Majiq, rMATS and orthogonal scores, respectively. The **gene_name**, **class_code**, **transcripfiname** and **exon_number** come from StringTie integration. The **class_code** matches how novel transcripts compare to annotated ones. The rMATS score column is missing from the image.

The main challenge for data integration with DJU method results is the difference in individual method implementation. For example, `JunctionSeq` does not produce splicing events. It computes the fold-change of introns, and because it adopts a distinct metric, it handles the effect size comparison with other tools inviable. DJU methods also use different definitions to define the coordinate of the AS events. Due to that, we have decided not to integrate effect-size, the PSI, from various methods, despite understanding this attribute is critical for alternative splicing identification.

This manuscript applies a benchmark to 2 independent datasets, the SIRV and the paired Illumina-ONT RNA-seq datasets. Other DJU method performance comparisons have used simulated datasets or datasets with a small subset of PCR-validated AS events[43, 9]. These 2 approaches do not fully appreciate the complexity of the RNA-seq experiment. Other benchmarks integrate splicing events on the gene level. Baltica can systematically resolve the different coordinate systems and compare multiple methods without these limitations.

The benchmark with the SIRV transcriptome is a special case that embeds a small complex transcriptome into a human transcriptome. The SIRV transcriptome has 138 introns, of which 2/3 change between the 3 contrasts. All SIRV introns are annotated, benefiting methods that rely on the transcriptome annotation, such as `JunctionSeq`. In addition, it contains almost 2:1 number positive to negative instances ratio, donor and acceptor splice with non-canonical sequence, and transcripts in opposite strands that share the identical introns. These attributes make the SIRV an important dataset for the benchmark of differential splicing identification methods.

In addition, we also benchmark the methods with the paired Illumina and ONT RNA-seq. Second-generation sequencing, primarily Illumina RNA-seq, has promoted many discoveries in differential AS. However, Illumina RNA-seq is limited by its relatively short read length, leading to a limited resolution of one single intron at a time. Third-generation RNA-seq technology, represented by Iso-seq from Pacific Biosciences (PacBio) and ONT RNA-seq, overcome this issue by offering longer sequencing reads than second-generation RNA-seq. The longer reads enable unambiguous matching to multiple introns in transcript isoforms and thus allow a better resolution of the transcriptome structure[44, 45]. Hybrid sequencing approaches pairing third-generation sequencing, and second-generation sequencing can benefit from both the deep coverage and the long-reads to improve AS identification task[46, 45, 47, 48]. This dataset allows us to compare the methods scores on 65,408 introns that have been tested for DJU with EdgeR in the ONT RNA-seq dataset.

We understand the Nanopore DJU scores are not a *bona fide* ground-truth, but only an orthogonal approach to the splicing identification problem. Although the apparent differences between the SIRV and the ONT RNA-seq datasets, the benchmark results were remarkably similar. Specifically, the ranks of DJU methods were the same in both datasets. On the other hand, the difference in F1 metric among methods was less pronounced in the second dataset.

In addition to the considerations detailed above, readers should interpret the benchmark results in the context of the 2 datasets. For example, the 2 benchmark use cases use transcriptomes with known introns, and they should benefit from methods that rely on a complete annotation. However, to a certain extent, current methods like `LeafCutter` and `Majiq` don’t rely on the annotation for the intron count modeling. Also, while `Majiq`, `LeafCutter`, and `rMATS` output events that comprise multiple SJ, `JunctionSeq` output a single *p*-value per intron. Moreover, for the second benchmark, one must keep in mind that EdgeR, which was used to analyze the Nanopore data, and `JunctionSeq` use a similar statistical model.

We demonstrate that the integration of DJU methods results can be helpful for intron prioritization. To do so, we have trained multiple machine learning models using 2 algorithms and five feature sets, one for each method score or the combination of the 4. We observed that models trained in the combined feature set performed better than methods trained in a single feature independent of machine algorithms. The LRC method has fewer parameters, requires fewer input data, and is faster to train but has lower predictive performance than GBC[49]. We used the LRC algorithm as a base model, intending to build upon it with the GCB. However, the 2 models presented a comparable performance. We provide the source code for model training and validation so that users can apply the method in their datasets. To our knowledge, this approach is the first method to take advantage of DJU scores to train a meta-classifier. However, this practice is common for bioinformatics practices and has been used to train predictors for protein secundary structure [50]. The integration helps prioritize introns for further experimental validation and may expand our knowledge on functional consequences of AS events.

We plan to extend Baltica in the future with additional DJU methods workflows and will include unit tests for tracking output differences due to changes in software versions. We invite the user community to follow the Baltica repository https://github.com/dieterich-lab/baltica for the code and update documentation and to participate in Baltica development.

## Key Points

- Methods to identify differential splicing changes are critical to detect the association of introns to genomic features such as genetic variants or splicing factor binding sites.
- However, these methods differ in many aspects, and the lack of standardization hampers the comparison of their results
- Baltica enables reproducible execution and integration of DJU methods. The integrated results allow benchmarking the different methods, and it reveals JunctionSeq ranks first in F1 metric across 2 independent benchmarks.
- Meta-classifiers trained on methods scores outperform all models trained in single method scores, demonstrating the data integration advantage.

## Acknowledgments

We are grateful to members of the Dieterich Lab for feedback on this work. Specifically, we would like to thank Tobias Jakobi and Harald Wilhelmi for the technical setup and feedback. The authors would like to thank Jennifer Gerbracht for sharing her dataset and her feedback on the earlier stages of the project. All authors acknowledge the support of the West-German Genome Center (WGGC) and, in particular, Tobias Lautwein.

## Author contributions

V.B. has tested the framework. V.B and N.G. have performed cell culture, library preparation, and sequencing. C.D. has processed the data. C.D. and N.G. supervised the project. T.B.B. processed data, developed and implemented the computational workflows and analysis scripts, tested the framework, led the analyses, contributed to the figures, and wrote the paper. All authors contributed to the research, including analysis, manuscript writing, and feedback for the framework design and development.

## Data availability

The SIRV RNA-seq is available at ArrayExpress E-MTAB-8461. The paired RNA-Seq and Nanopore-Seq are available from CD and NG upon reasonable request. In addition, data matrices used for the benchmark and meta-prediction are available at https://doi.org/10.5281/zenodo.5643428.

## Funding

This work has been supported by Informatics for Life, funded by the Klaus Tschira Foundation and the Deutsche Forschungsgemeinschaft (DFG, DI1501/8-2, and GE2014/6-2). Nanopore sequencing was supported by a DFG Sequencing Call grant to CD (DI1501/12-1, project 423957469).

**Thiago Britto-Borges** is a postdoctoral researcher in Biomedical Data Science at the University Hospital Heidelberg.

**Volker Boehm** is a postdoctoral researcher in RNA Biology at the Institute for Genetics, University of Cologne.

**Niels Gehring** is a professor at the Institute for Genetics, University of Cologne.

**Christoph Dieterich** is a full professor and head of the Klaus Tschira Institute for Integrative Computational Cardiology at the University Hospital Heidelberg.

## Supplementary Materials

Here we provide the supplementary information to the manuscript Baltica: integrated splice junction usage analysis, by Thiago Britto-Borges, Volker Boehm, Niels H. Gehring and Christoph Dieterich.

### Experimental design for the SIRV dataset

**Table S1.**
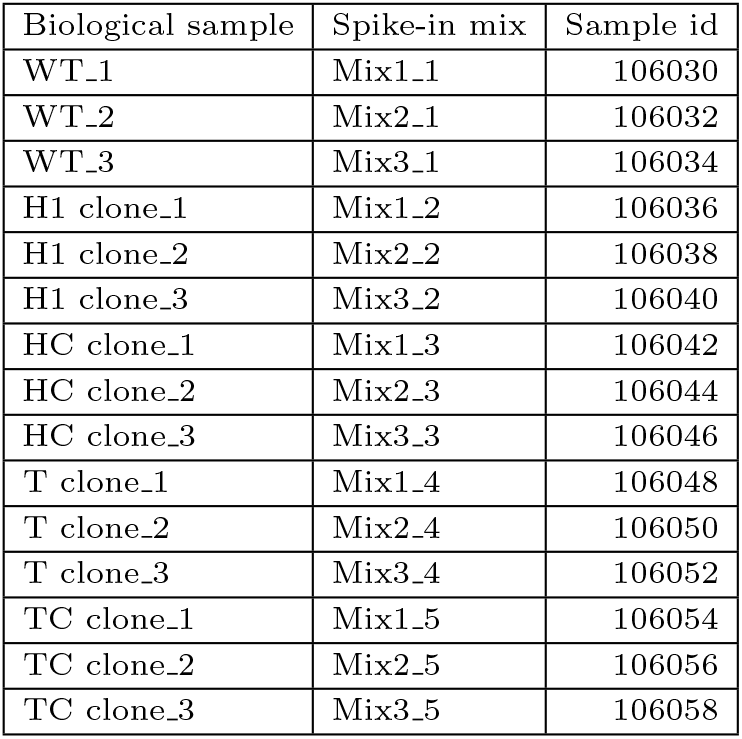
The biological sample, spike-in mix, and sample identifier associations for samples in dataset E-MTAB-8461. Related to Figure 2. Note that the biological samples and the spike-in mixes are not confounded.

### Popularity of DJU methods

**Fig. S1.**
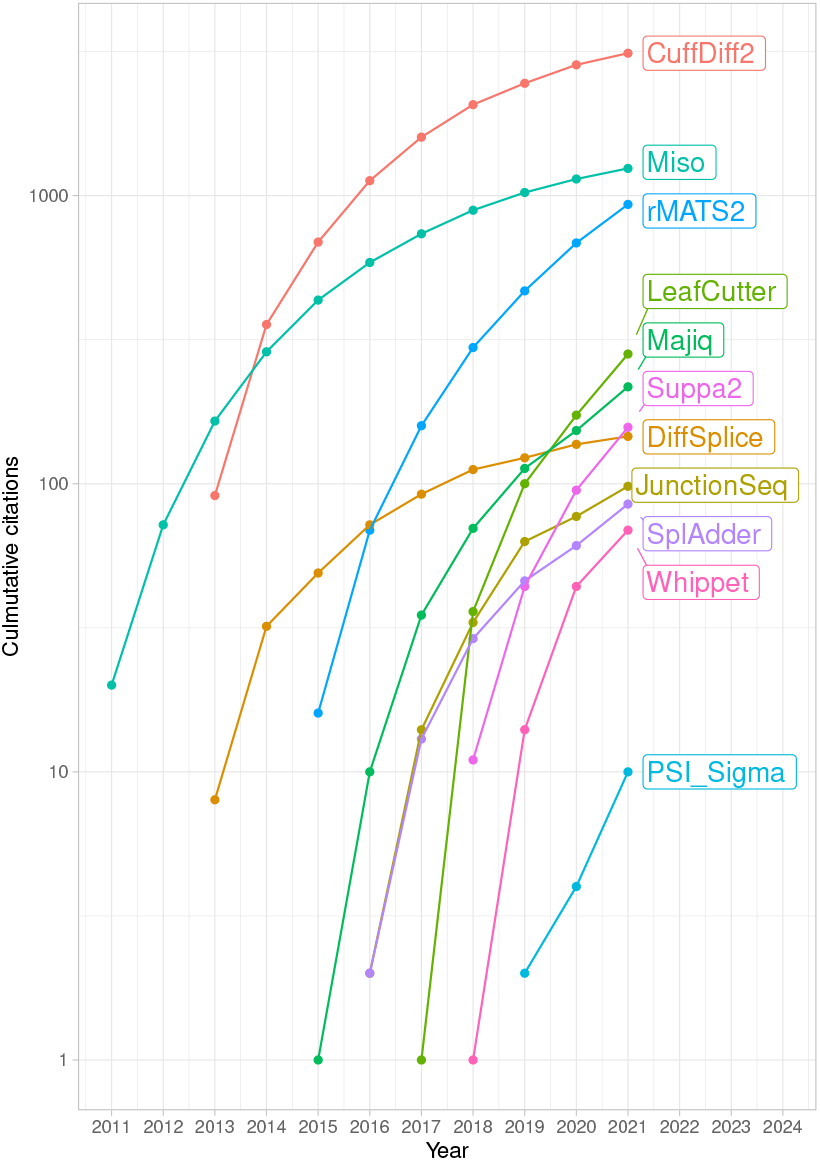
The popularity of selected methods for differential splicing identification. The citation over time for selected DJU methods. The y-axis is *logio* transformed. Data sourced from https://scholar.google.com/ using the scholar R package.

### Heatmap for the SIRV benchmark

**Fig. S2.**
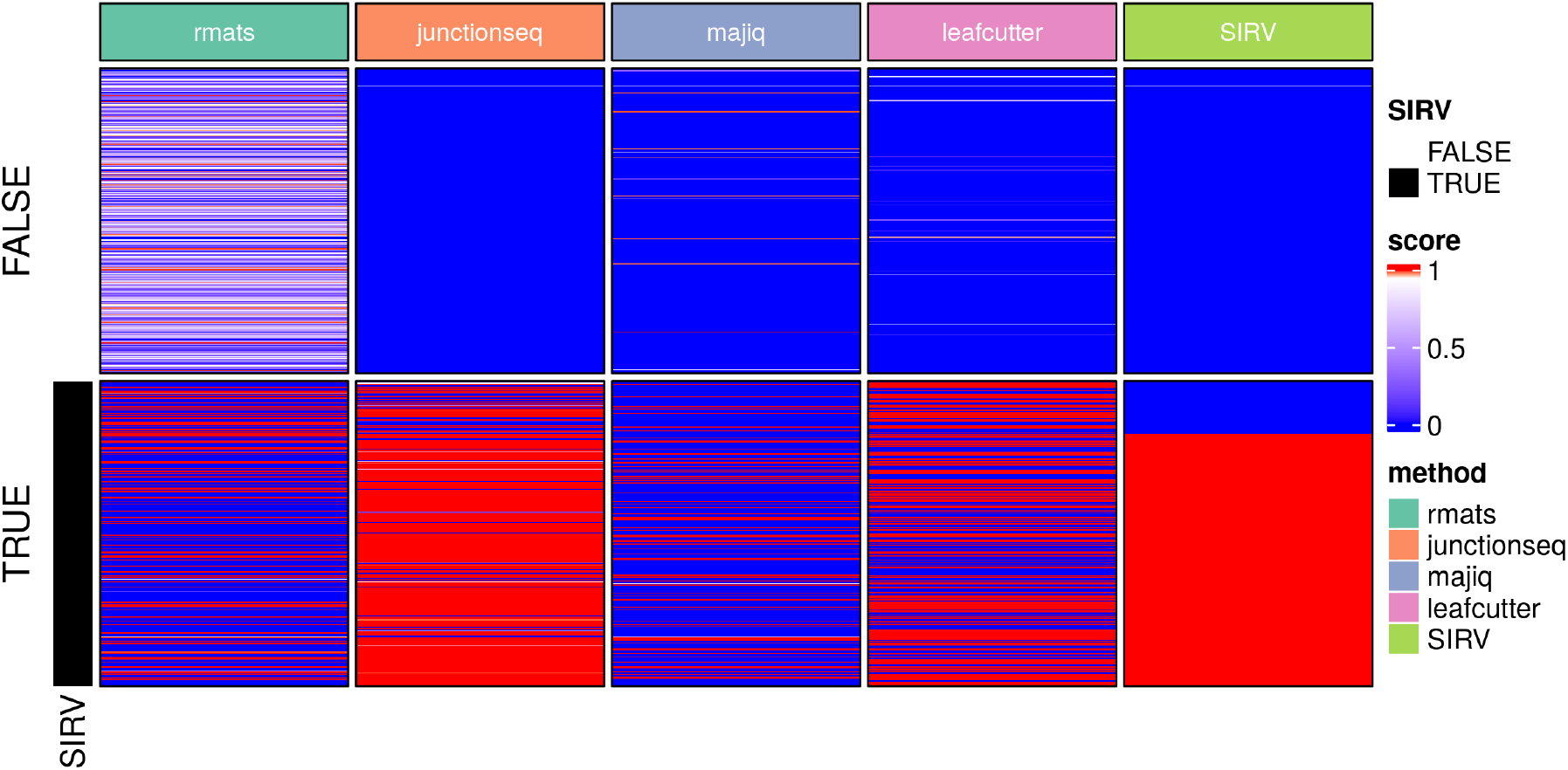
Four hundred fourteen introns were randomly sampled from the human transcriptome (top) and the same amount in the SIRV transcriptome (bottom). Introns in the SIRV transcriptome are annotated, as shown in the leftmost annotation. Colorbar shows method score as divergent color from red (1) to blue (0), and the highest score better the performance classification. The methods score for negative instances, in the top panel, agreement while for positive, for instance, the 2/3 in the bottom panel, are not.

### SIRV benchmark precision-recall curve

**Fig. S3.**
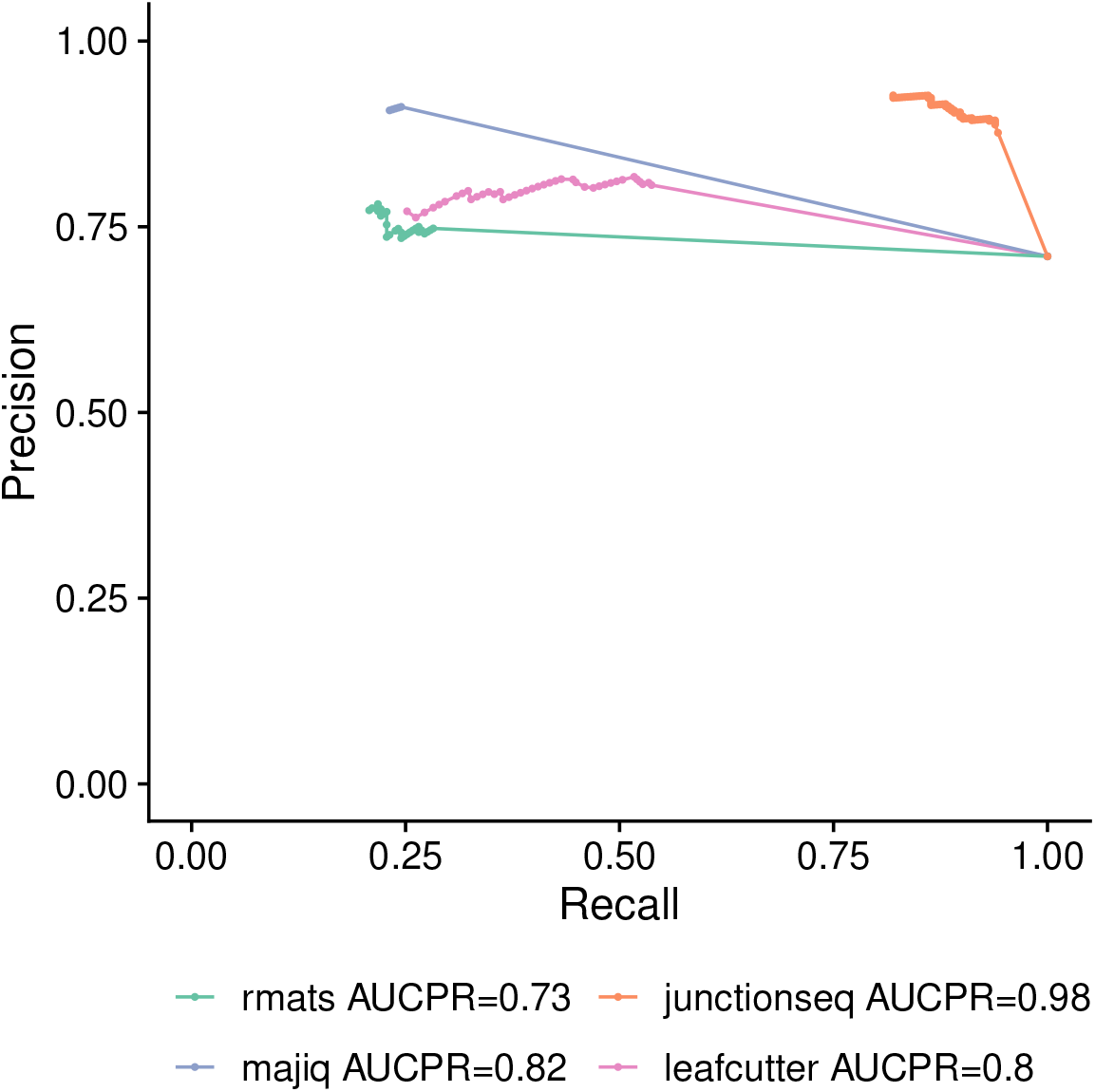
SIRV benchmark precision-recall curve. Overall, changes in recall have only a slight effect on precision for all the methods. `JunctionSeq` ranks first (AUCPR=0.98), followed by `Majiq` (AUCPR=0.82), `LeafCutter` (AUCPR=0.8), and `rMATS` (AUCPR=0.73). PR: precion-recall; AUCPR: area under the precision-recall curve. Relates to Figure 3.

### Spearman correlation for the SIRV benchmark

**Fig. S4.**
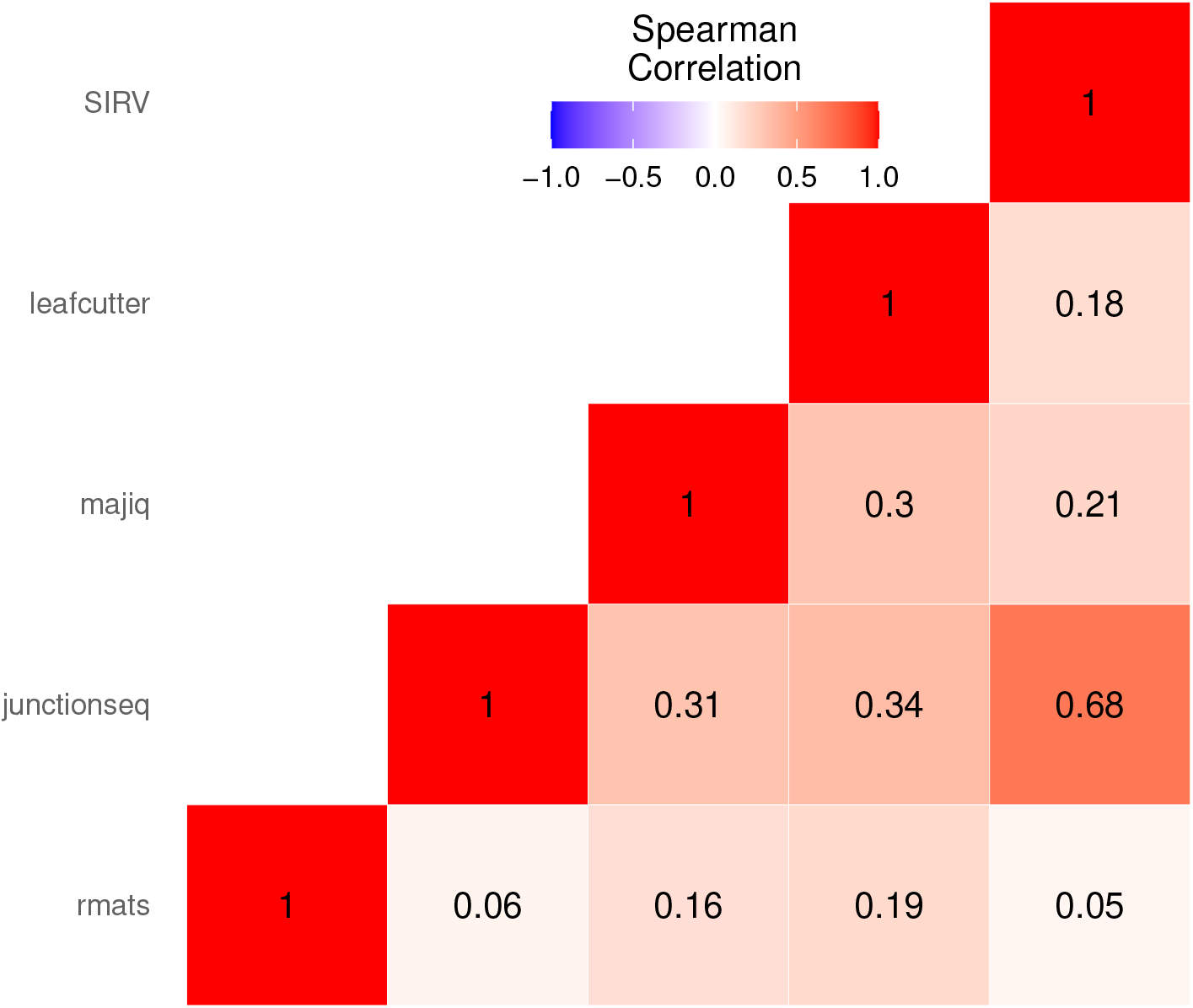
Spearman correlation among methods and the SIRV ground-truth. Scores were log transformed with the function *f*(*x*) = −*log*_110_(*x* + 1*e*^−10^) and only introns from the SIRV transcriptome were tested. Overall, pairs of methods have a low correlation with an mean of 0.25.The maximum value is of 0.68 for l and ground-truth.

### Perfomance metrics and confusion matrix for the SIRV benchmark

Metrics were obtained with caret::confusionMatrix function. True and false instances were separate with the 0.95 threshold.

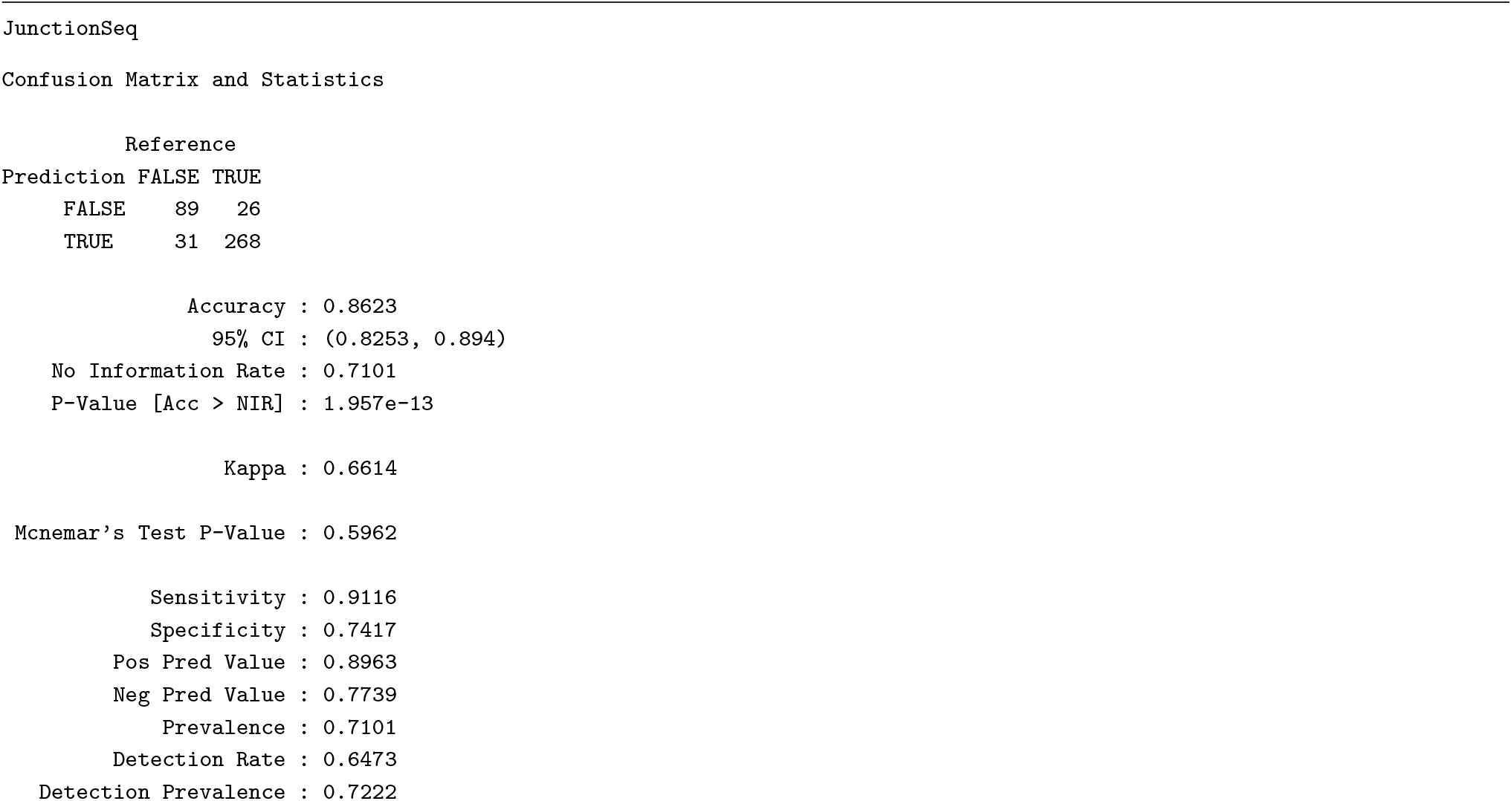
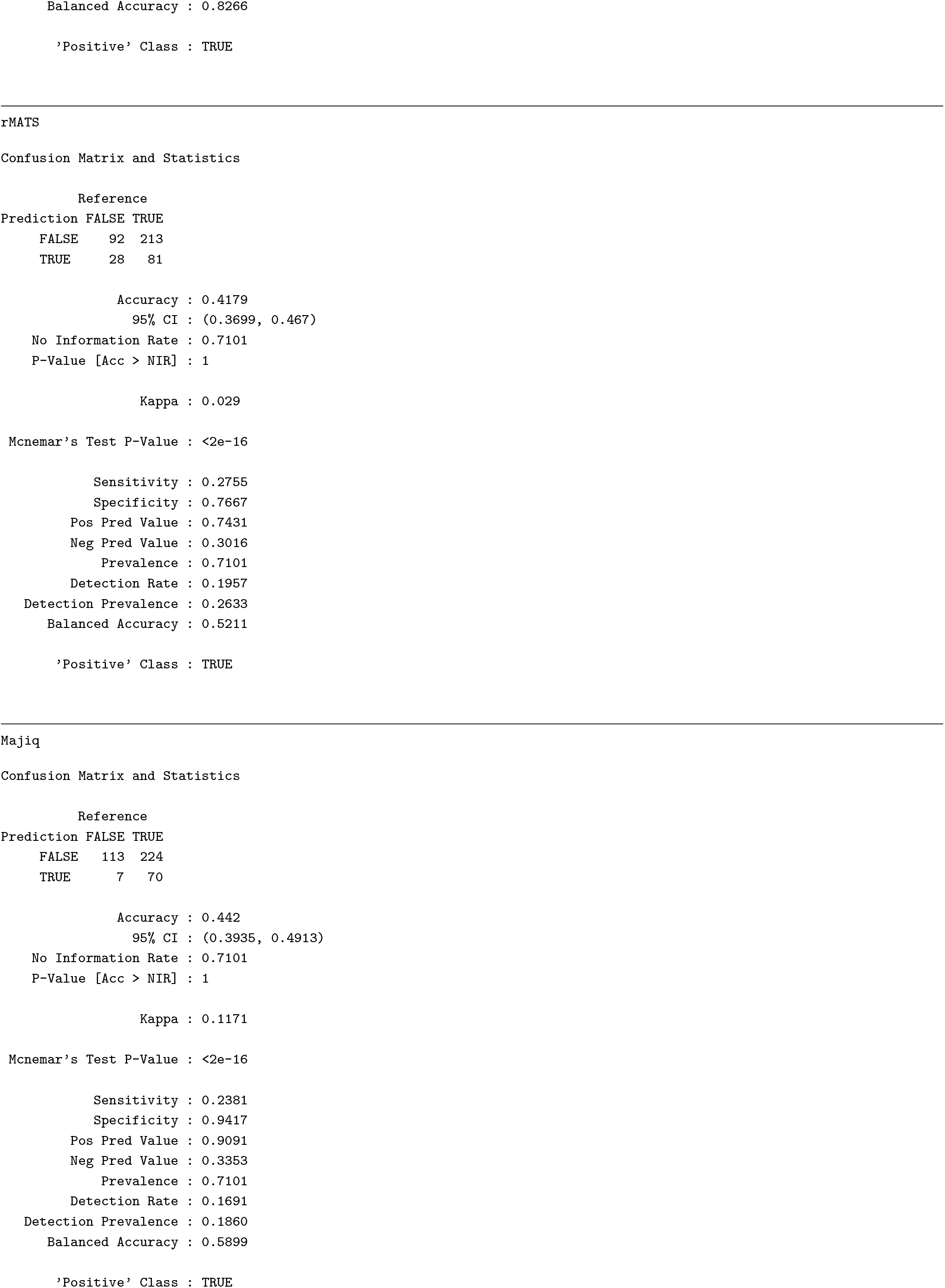
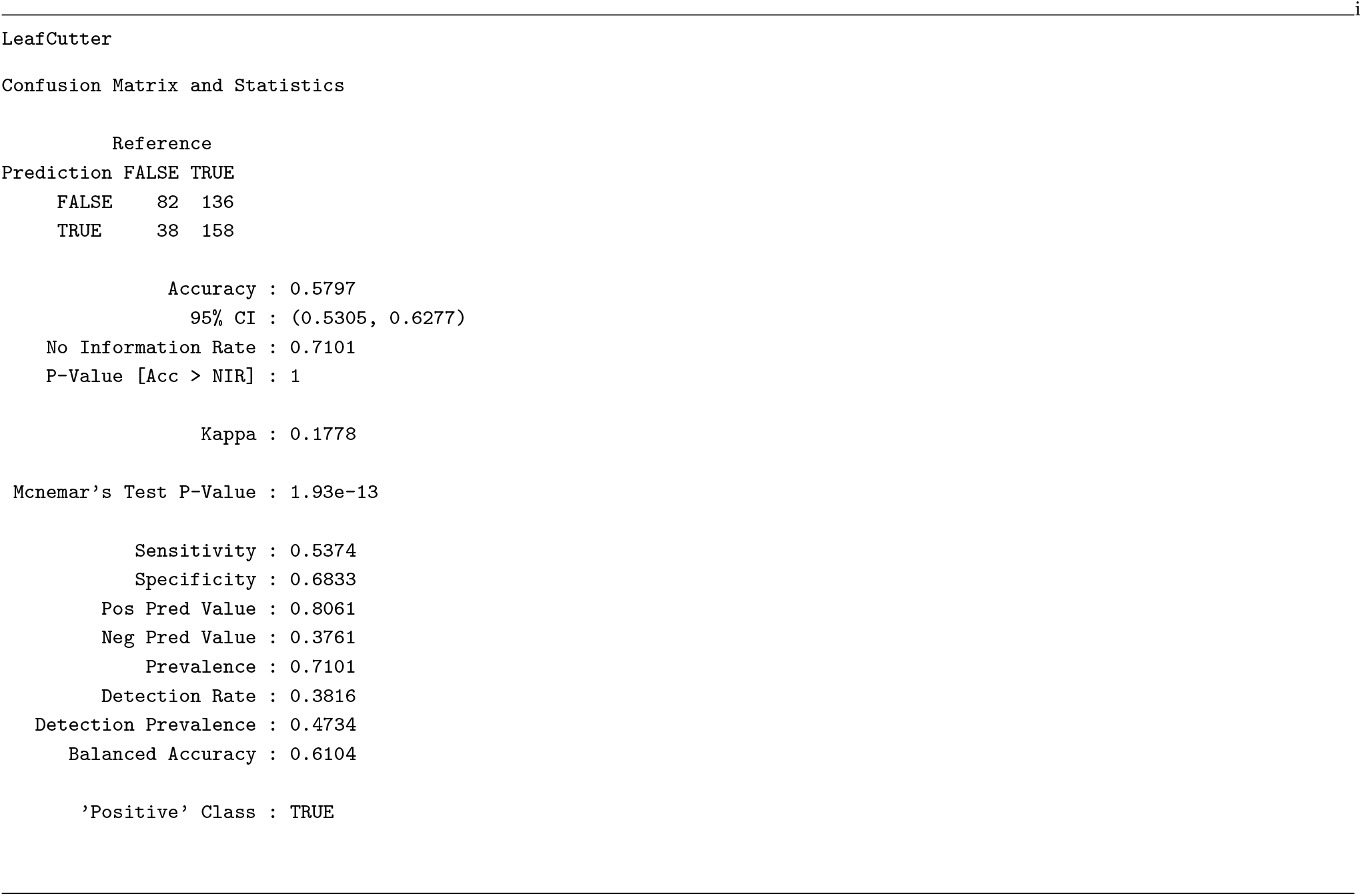

### Baltica integration for the ONT RNA-seq dataset

**Fig. S5.**
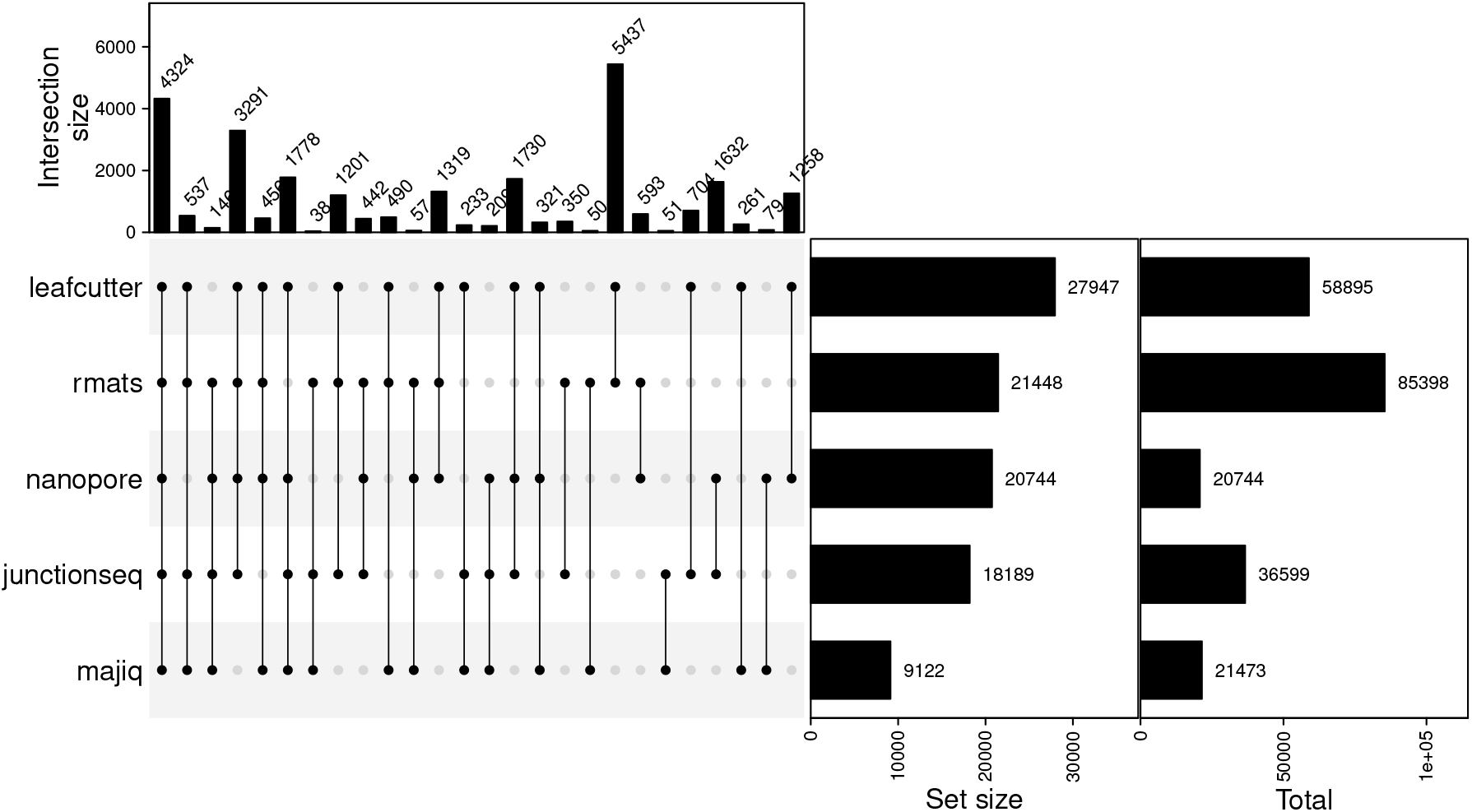
Baltica integrated SJ for the ONT dataset. The plot shows distinct sets of introns with *score* > 0.95 by combinations of methods and the DJU for the ONT RNA-seq. The complement sets, combinations with a degree of 1, were omitted. The intersection and set sizes refer to the number of the sets not omitted, while the total shows the total number of calls.

### ONT RNA-seq benchmark precision-recall curve

**Fig. S6.**
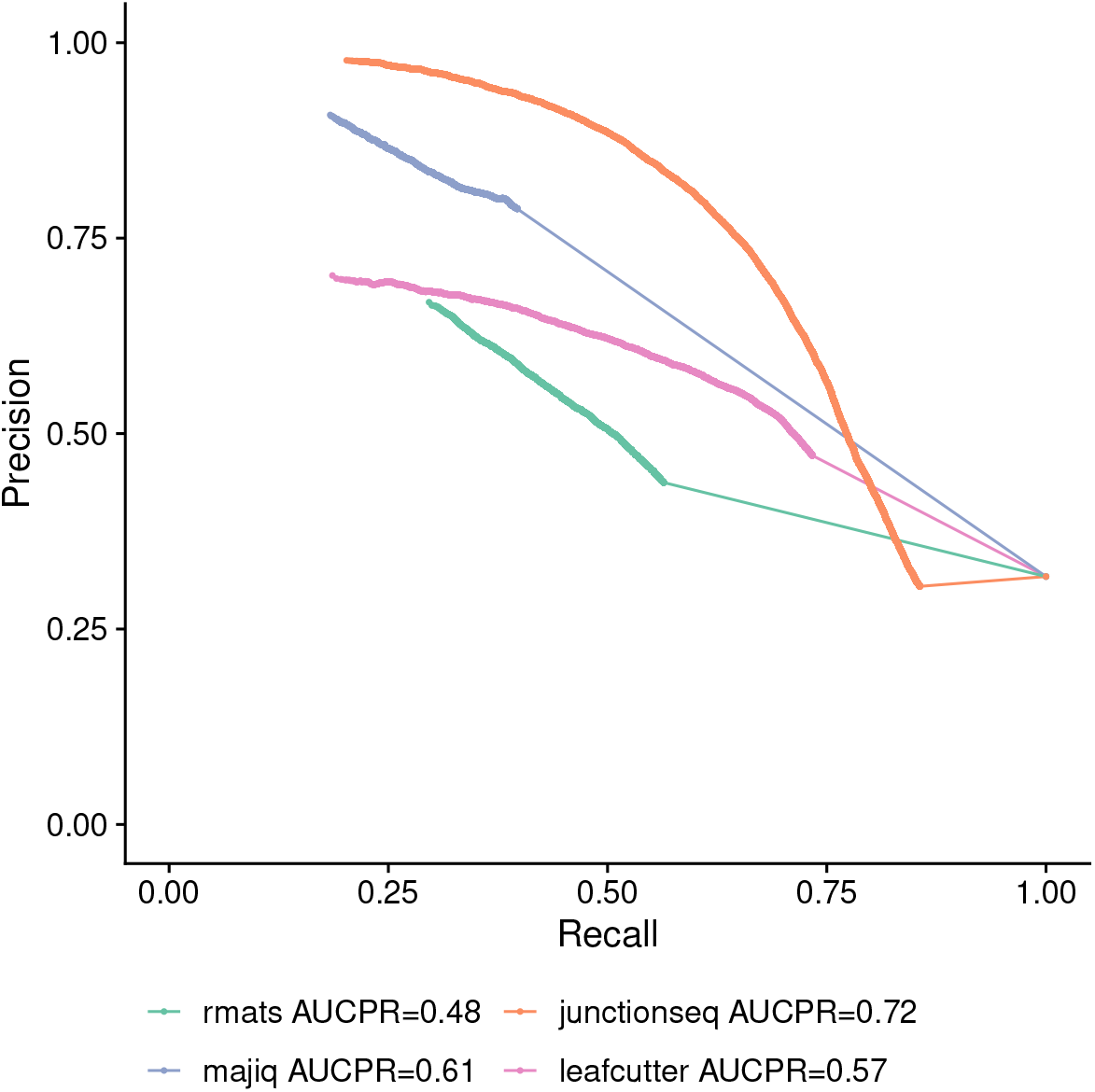
ONT RNA-seq benchmark precision-recall curve. Different from the SIRV benchmark, the precision-recall curve for the ONT RNA-seq benchmark shows negative correlation between precision and recall. However, the method rank is consistent with the SIRV benchmark with `JunctionSeq` first (AUCPR=0.72), and then `Majiq` (AUCPR=0.61), `LeafCutter` (AUCPR=0.57), and `rMATS` (AUCPR=0.48). PR: precion-recall; AUCPR: area under the precision-recall curve. Relates to Figure 4

### Heatmap for benchmark with the ONT RNA-seq dataset

**Fig. S7.**
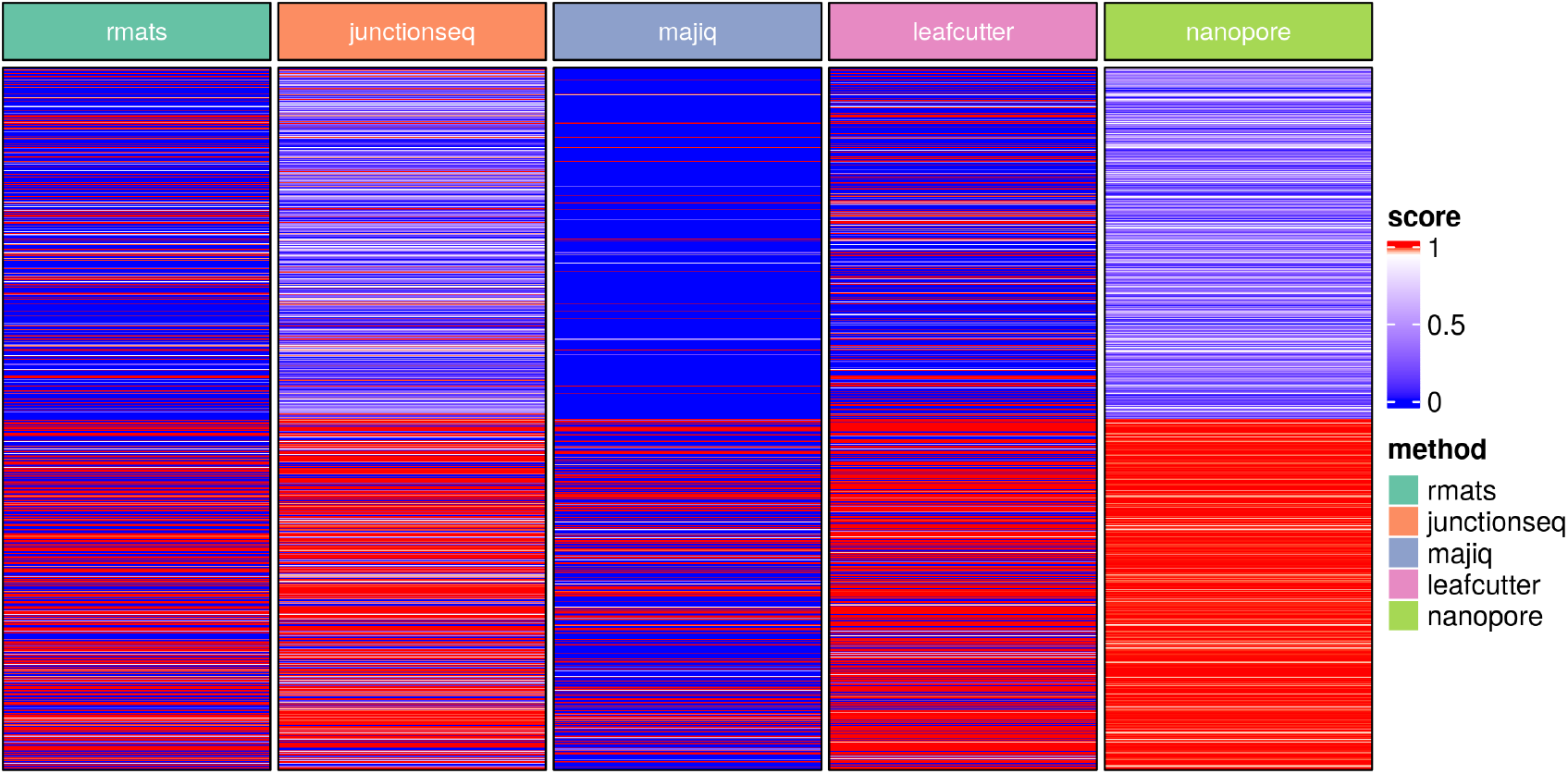
The 0.95 thresholds separate five hundred negative (top) and positive instances (bottom) detected by the ONT RNA-seq. Colorbar shows method score as divergent color from red (1) to blue (0), higher is better. Related with Supplementary Figure S2

### Spearman correlation among methods scores and ONT RNA-seq DJU method for the ONT RNA-seq dataset

**Fig. S8.**
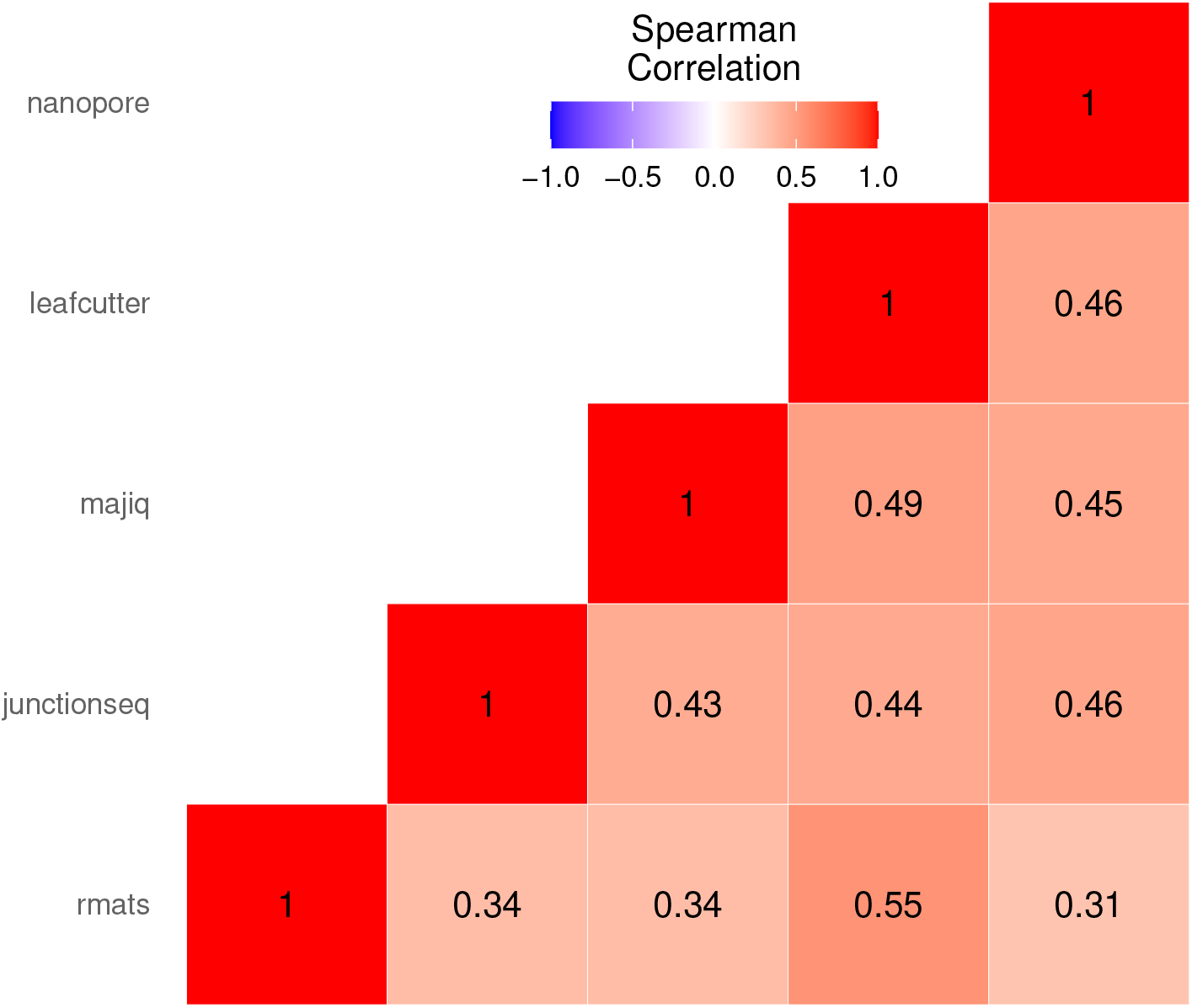
Spearman correlation among scores from DJU methods and ONT RNA-seq dataset DJU. Overall, methods scores show higher correlation than in the SIRV benchmark (Supplementary Figure S4), however the agreement between to methods is still limited, whith the maximum value of 0.55 for `rMATS` versus `LeafCutter`. Scores were transformed with *f*(*x*) = −*log*_10_(*x* + 1*e*^−10^) and only introns presented in the ONT RNA-seq dataset were tested.

### Confusion matrix for the benchmark with the ONT dataset

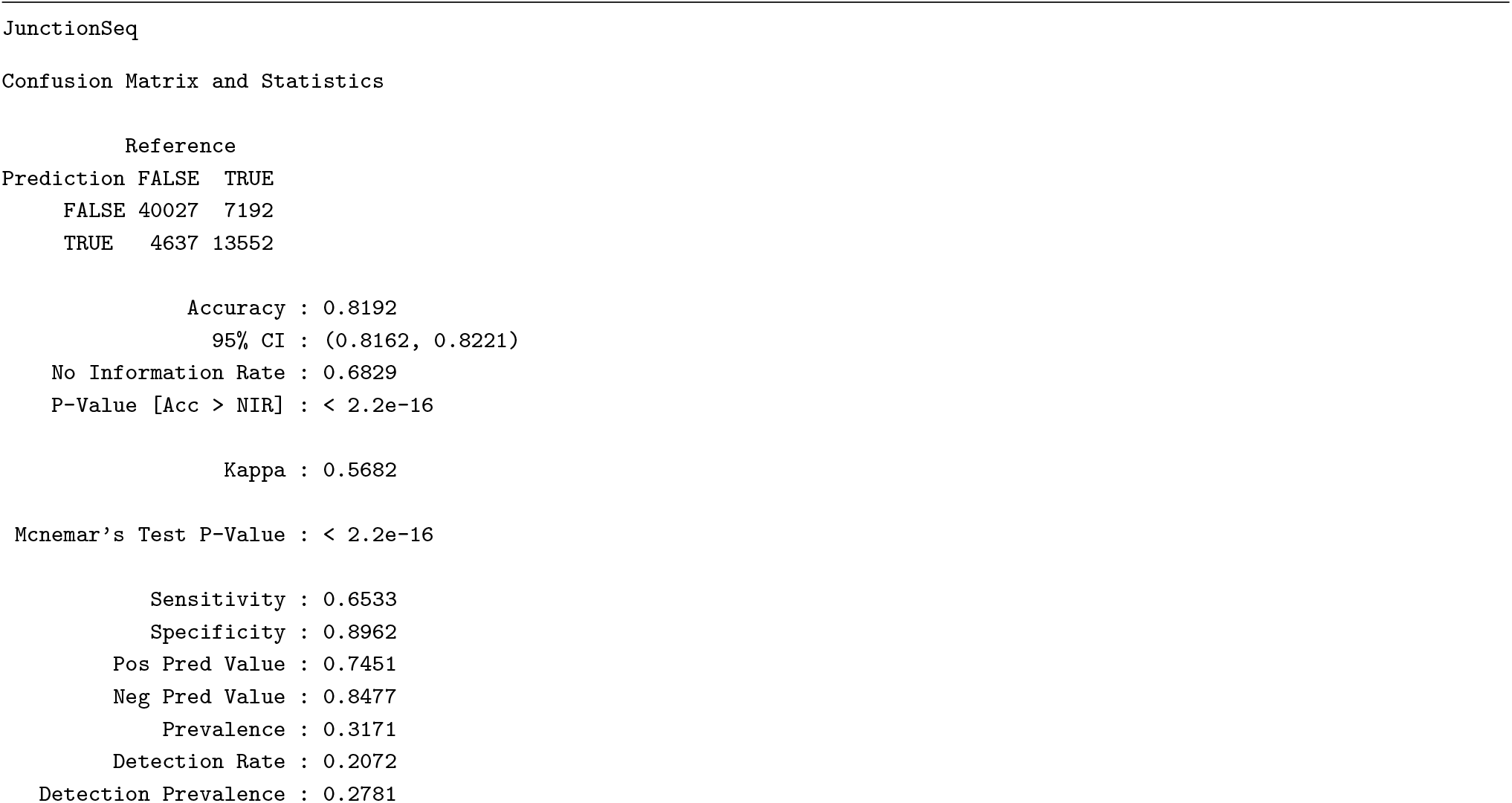
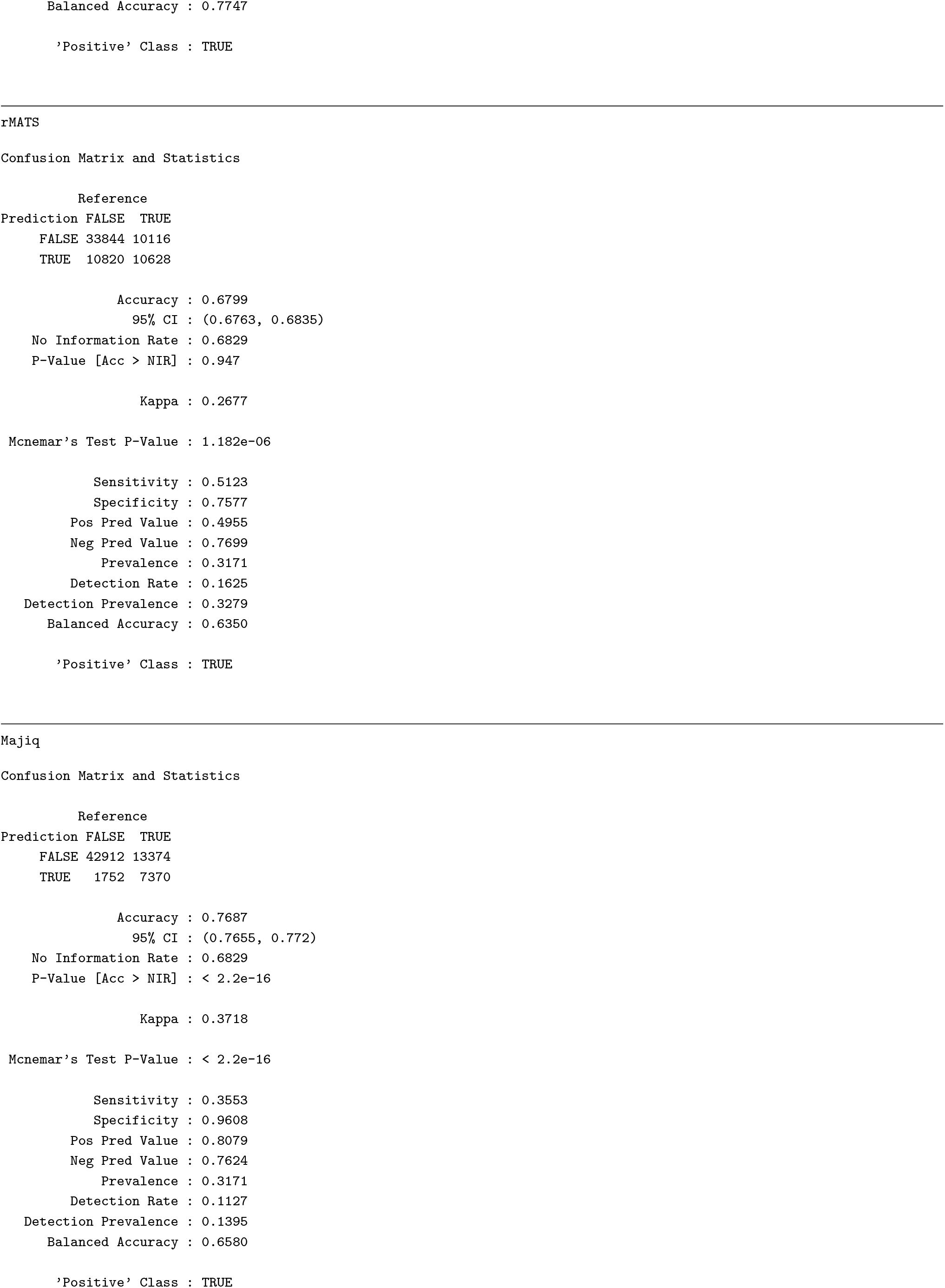
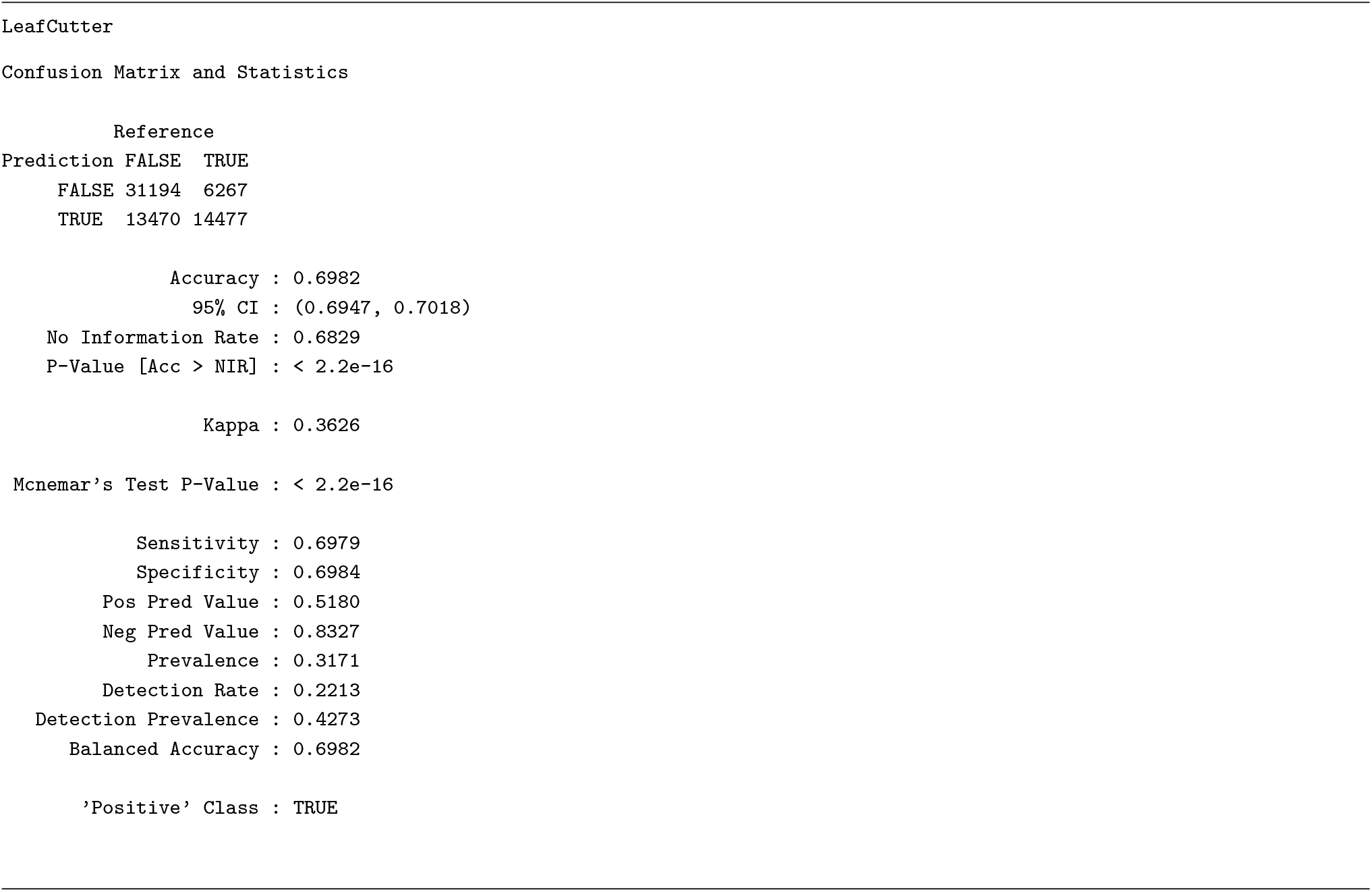

